# Atlas of Proteomic Technologies: an evidence-based framework for selecting and combining commercial proteomics platforms

**DOI:** 10.64898/2026.09.10.750723

**Authors:** Christopher D. Whelan, Karl Smith-Byrne

## Abstract

High-throughput proteomics now spans several technologies that differ in biological breadth, precision, specificity, sensitivity, cost, and method(s) of quantification; however, investigators currently lack an evidence-based framework to select or combine platforms. Here, we present the Atlas of Proteomic Technologies (APT) - a curated resource that evaluates six commercial platforms across ten analytical dimensions using published head-to-head evidence, expert evaluation, and iterative peer review. APT hosts two decision tools: the *Help Me Choose* tool maps a study’s primary aim(s), scale, sample matrix, and technical constraints to a calibrated, weight-adjustable platform ranking, and the *Help Me Combine* tool ranks complementary platform pairs by their net-new protein coverage and user-specified technical priorities. Both tools were tested exhaustively across all possible combinations and behave in a balanced, merit-based manner. APT is openly accessible at https://aptatlas.org, providing downloadable tool specifications, seeded analysis scripts, and complete protein coverage lists, ensuring every score and recommendation is inspectable and reproducible.

## Introduction

Proteomics and genomics are increasingly combined at scale to identify associations between protein concentrations and the risk(s) of disease endpoints and other health-related traits. Genetic studies of protein levels^1,2^ allow for causal inference tools like Mendelian randomisation and colocalization to uncover novel therapeutic targets^3^, while machine learning models trained directly on large-scale proteomic data can identify predictive biomarkers for dozens of diseases with unmet needs^4^. This expanding subfield of large-scale molecular epidemiology, commonly known as ‘population proteomics’ or ‘proteogenomics’, has transformed multiplex protein measurement from a specialized technology into a fundamental tool for population health researchers, clinical scientists, drug developers, and translational geneticists. In parallel, the technologies that enable population proteomics have proliferated, with multiple affinity platforms based on antibodies^5,6,7^ and aptamers^8^ now sitting alongside mass spectrometry (MS) workflows leveraging nanoparticle enrichment^9^ or data-independent acquisition (DIA) with hyper-reaction monitoring (HRM)^10^ to enable deep characterization of the human proteome at unprecedented scale and throughput.

The growing proteomic technological landscape and its diversifying user base present challenges for study design and data interpretation for research scientists. Certain technologies maximise the number of proteins measured (‘coverage’); others augment confidence that a signal reflects the intended target (‘specificity’); others still prioritise reliable measurements over time (‘precision’), detection of very low-expressing proteins (‘sensitivity’), absolute concentration output (pg/mL), or low cost per sample. A broad aptamer or MS assay may detect many thousands of proteins but at reduced specificity or reproducibility, while a targeted proximity panel may measure only a few hundred analytes yet resolve them at attomolar sensitivity. Because no single platform is optimized on every axis, the platform an investigator selects is a consequential early design decision, shaping which proteins are observable, whether findings will replicate across cohorts, how protein quantitative trait loci (pQTLs) can be interpreted, and how far a fixed budget can extend. Researchers without formal training or extensive experience in proteomics may struggle to navigate this increasingly complex subfield. Conversely, experienced labs or large industry consortia aiming to deploy two platforms in parallel to widen proteome coverage or cross-validate signals will face a distinct set of challenges. A pair of platforms can be highly complementary or largely redundant depending on how much net-new content the second one contributes, how their measurement classes balance, and how their combined cost and sample volume demands aggregate. Two broad platforms with similar catalogues may overlap so heavily that pairing them contributes little, whereas a broad platform paired with an orthogonal technology could extend coverage of proteins and pathways of interest substantially. Thus, pairing decisions are as consequential as single-platform decisions, but comparatively little data exist to support them.

The published literature contains several informative head-to-head benchmarking studies to support single platform decisions, mostly focusing on the two widely deployed affinity platforms: Somalogic and Olink. These studies have concluded that the platforms agree only modestly at the level of individual proteins, with median cross-platform correlations of approximately 0.3 to 0.6 between individual proteins and a consistently bimodal pattern in which a minority of proteins correlate strongly while 40-50% of proteins correlate poorly^11,12,13^. Genetic anchoring via pQTLs has been proposed as an arbiter of assay quality, with dual antibody-based Olink assays generally yielding more *cis*-pQTL support than aptamer-based SomaScan assays; however, both platforms carry comparable rates of epitope artifacts, that is, genetic variants that alter the protein epitope recognized by the antibody or aptamer and shift the measured signal independently of true protein abundance^14,15^. Meanwhile, MS-based platforms access a largely distinct, higher-abundance fraction of the proteome^16^, contributing dozens of otherwise undetected proteins robust to epitope artifacts. Thus, most comparative technological evaluations have concluded that antibody-, aptamer-, and MS-based technologies are complementary rather than interchangeable, with no single platform dominating across coverage, precision, sensitivity, and quantification^16,17,18^. Nonetheless, these head-to-head studies compared only subsets of the broader technological landscape using specific cohorts and limited endpoints, reporting snapshots of platform performance that age as products iterate and may insufficiently address specific scientific use cases. A study optimised for pQTL discovery could reach different conclusions from one optimised for clinical biomarker validation, and neither may be appropriately designed to study biological subtypes in post-mortem brain tissues. The field currently lacks a resource that consolidates the comparative evidence into a common, transparent framework; lets investigators set relative weights for the dimensions that matter for their own study; and facilitates decision-making both for single platforms and combined implementation, without collapsing everything into a single ranking that hides the assumptions behind it.

We built the Atlas of Proteomic Technologies (APT) to address this gap. APT is an open web resource that curates comparative evidence for six commercially available proteomics platforms and scores each on ten analytical dimensions, recording the supporting citation and reasoning for every score. APT’s evidence base facilitates two decision tools. The first tool, ‘Help Me Choose’, converts a study’s primary goal, sample matrix, scale, and mandatory requirements into effective dimension weights and returns a calibrated platform ranking, with each recommendation contextualized by the inputs that drove it. The second tool, ‘Help Me Combine’, extends the same framework to platform pairs, ranking the fifteen combinations of proteomics platforms by how much net-new information each pair adds under the priorities the user selects. Here, we describe the construction of APT and its underlying scoring system, the development and iterative revisions of its single-platform recommendation tool, the proteome coverage analyses that underpin its platform combination tool, and how both tools were tested for fairness across up to 1.2 million permutations to help minimize biases and produce data-driven, context-dependent results.

## Results

### Platform Inclusion

Proteomics platforms were included if they were commercially available and actively marketed for biomedical research, measured at least 100 proteins in at least one blood-based matrix, and had at least one peer-reviewed publication in a human cohort. Where a technology had evolved across generations, we scored the most current configuration (STAR Methods, Platform selection). Pre-commercial, legacy, and single-analyte assays were excluded. From an initial set of 46 surveyed protein measurement platforms (**Supplementary Table S1**), our criteria identified six commercially available plasma proteomics platforms for further evaluation. The six platforms included Olink Explore HT (proximity extension assay), Illumina SomaSeq Discovery (hereafter SomaSeq; the next-generation-sequencing SOMAmer / aptamer readout and successor to the array-based SomaScan assay), Alamar NULISA (a sensitive proximity ligation panel), Nomic nELISA / Omni 1000 (a bead-based sandwich immunoassay), Seer Proteograph XT (DIA-MS with nanoparticle enrichment), and Biognosys TrueDiscovery (DIA-MS with HRM) (**Table 1**). Together, these technologies cover the main measurement classes now in routine use, from single-binder and dual-binder affinity assays to enrichment-based and direct DIA MS. **Figure 1** summarises the pipeline we followed to build this resource and the tools it operates through.

**Figure 1.**
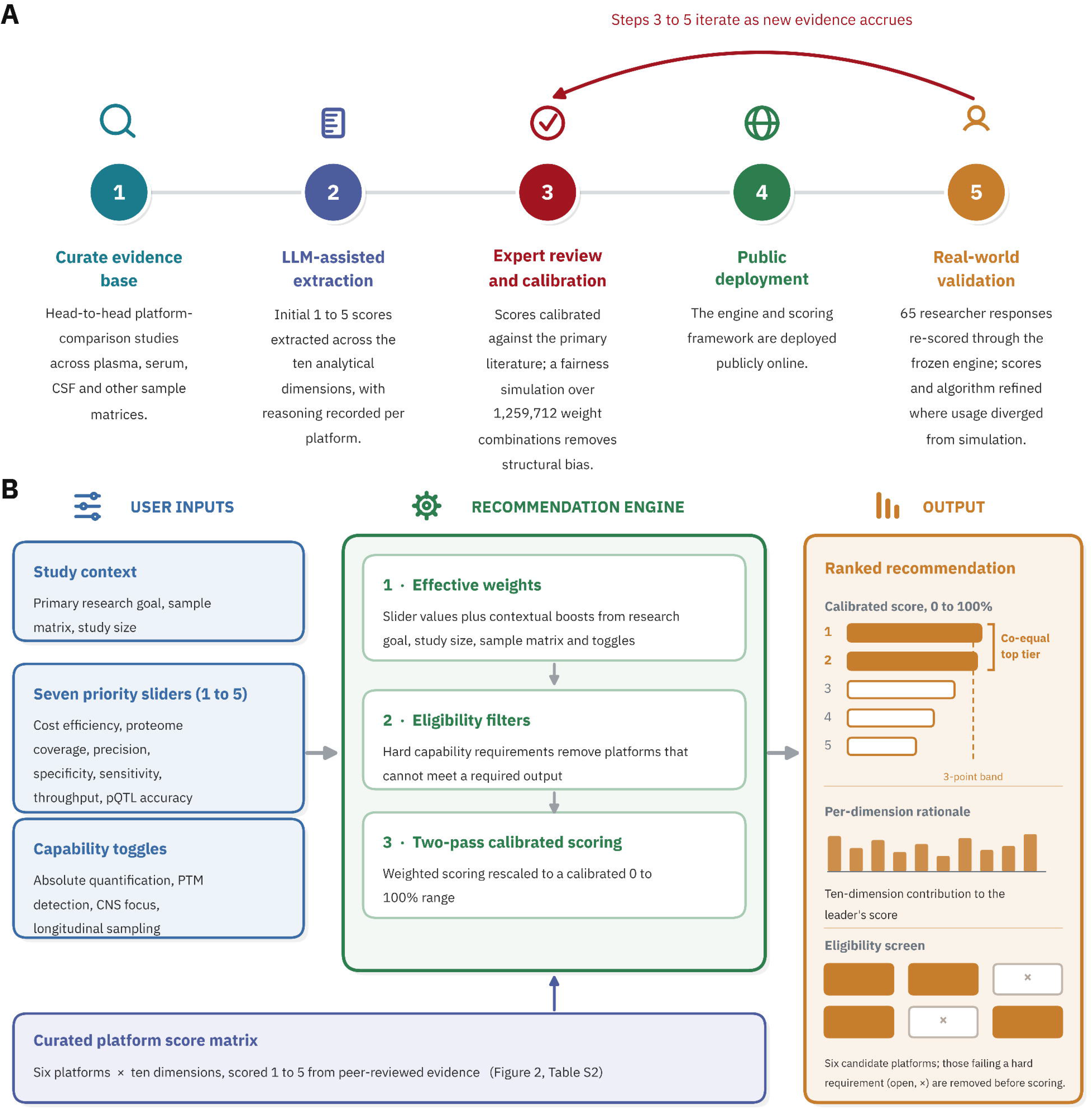
Development and operation of the Help Me Choose tool. (A) Five-stage development pipeline. As an initial step (#1), we curated a collection of head-to-head platform comparison studies and fed them to a large language model (#2; Claude Opus 4.8) to compute initial 1-to-5 scores across ten analytical dimensions, recording its reasoning for each platform. We then manually calibrated these scores against the primary literature (#3), updating them based on peer-reviewed evidence and expert assessments. We next ran a fairness simulation over 1.26 million weight combinations to remove structural bias, followed by public deployment (#4) and real-world validation (#5) in which 63 researcher responses were re-scored through the frozen, production-version tool. Steps 3 to 5 will iterate as new evidence accrues, guiding revised scoring in future iterations of the app. (B) Operating schematic. User inputs (study context, seven priority sliders, and capability toggles), together with the curated six-platform by ten-dimension score matrix (Figure 2**, Supplementary Table S3**), feed a three-stage tool incorporating effective weights, eligibility filters, and two-pass calibrated scoring. Platforms are ranked by their calibrated scores (0% to 100%), with those within a 3-point band of the leader reported as a co-equal top tier, alongside a per-dimension contribution breakdown and an eligibility screen.

**Table 1.** Summary of the six evaluated plasma proteomics platforms. Approximate protein counts reflect current commercial configurations. These vary by product version and, for mass spectrometry, by study and spectral library. Full dimension scores (1 to 5) with supporting citations are provided in Supplementary Tables S3 to S9. DIA = data-independent acquisition. HRM = hyper reaction monitoring. iRT = indexed retention time. MS = mass spectrometry. NPX = normalised protein expression. RFU = relative fluorescence units. NPQ = NULISA protein quantification.

| Platform | Technology and chemistry | Readout and front end | Approximate number of proteins | Quantification | Notable strengths |
| --- | --- | --- | --- | --- | --- |
| Olink Explore HT | Proximity extension assay (dual antibody) | Next-generation sequencing (NGS); liquid handler + thermal cyclers on front end | 5,400 | Relative (NPX) | Throughput (5/5), specificity (4/5) |
| Illumina SomaSeq Discovery | Aptamer (SOMAmer) | NGS; SomaSeq automation system on front end | 9,500 | Relative (RFU) | Proteome coverage (5/5), precision (5/5) |
| Alamar NULISA | Proximity ligation assay (dual antibody) with assay background suppression | NGS; ARGO HT instrument on front end | 220 (Neuro 220)250 (Inflammation 250) | Relative (NPQ) | Sensitivity (5/5), neuroscience-relevant PTMs (e.g., p-tau217) |
| Nomic nELISA Omni 1000 | Bead-based sandwich immunoassay with encoded microparticles | Flow cytometry; four-laser cytometer + automated liquid handling on front end (service only) | 1,058 | Absolute - referenced (pg/mL) | Cost per sample (5/5), quantification (4/5) |
| Seer Proteograph XT | DIA MS with nanoparticle enrichment | Mass spectrometry; SP100 instrument + LC-MS on front end | 8,500 to 10,700 (instrument-dependent) | Relative (label-free intensity) | Coverage (5/5), pQTL accuracy (5/5) |
| Biognosys TrueDiscovery | DIA MS (HRM, iRT-referenced) | Mass spectrometry; automated depletion/digestion + LC-MS on front end (service only) | 3,575 to 7,000 (instrument- and P2 enrichment-dependent) | Relative (label-free intensity) | Multi-matrix validation (5/5), specificity (4/5) |

### Scoring Dimensions

Each platform was scored on an integer scale from 1 (poor relative to the current field) to 5 (best-in-class) across ten dimensions, including (1) proteome coverage, (2) measurement precision, (3) target specificity, (4) sensitivity, (5) cost per sample, (6) throughput, (7) quantification type, (8) multi-matrix validation, (9) evidence depth, (10) and pQTL accuracy. The scores are deliberately relative, expressing where a platform stands within the evaluated set, and not against an absolute external standard. For example, a score of 3 represents average for the field. Full dimension definitions and scoring criteria are provided in **Supplementary Table S2**, and every platform score is accompanied by its supporting citation(s) and rationale in **Supplementary Tables S3 to S9** (summarised in **Table 1** and **Figure 2**).

**Figure 2.**
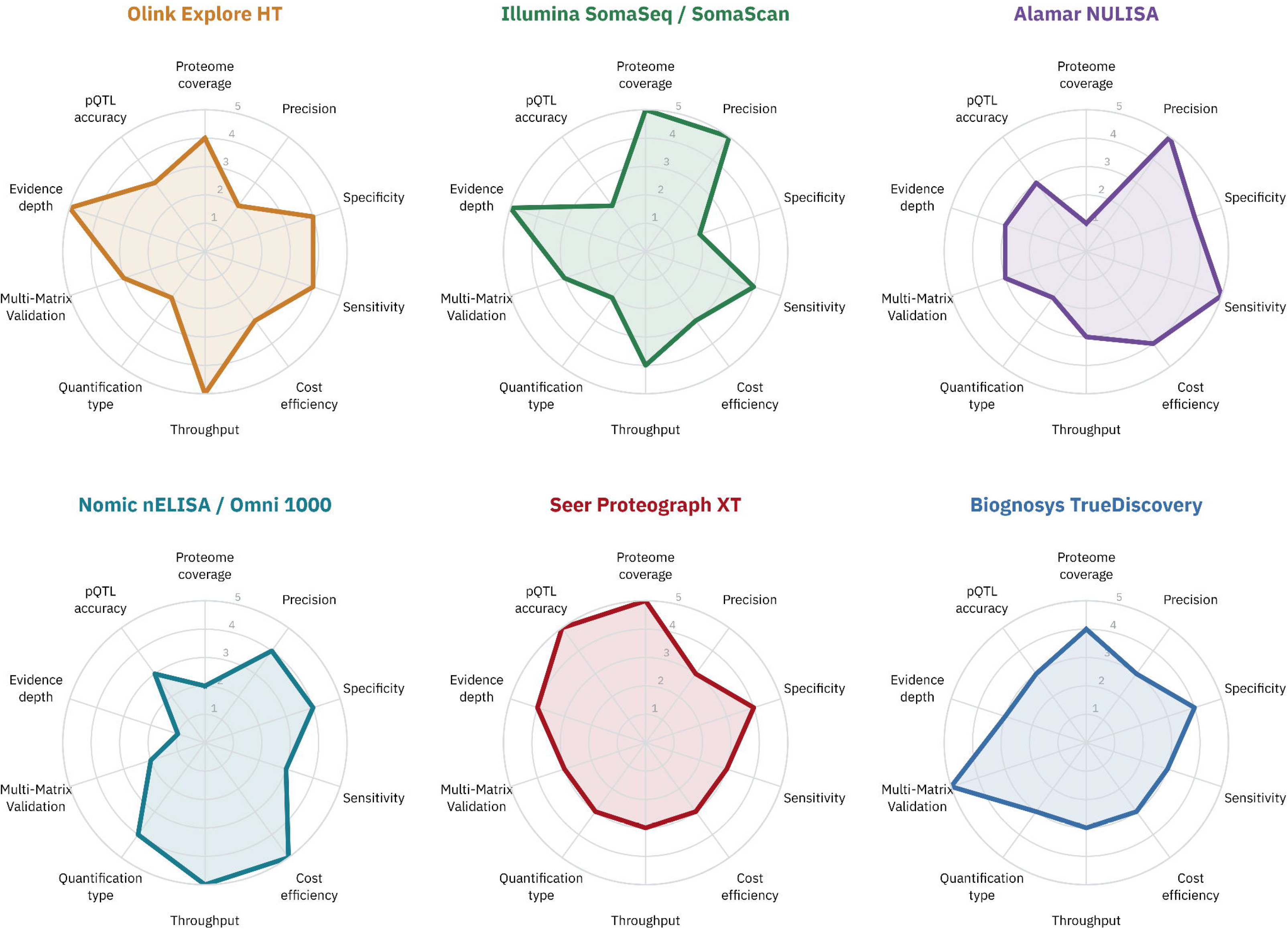
Platform score profiles across the ten analytical dimensions. Radar plots of the six evaluated platforms, each scored from 1 (centre) to 5 (outer ring) on proteome coverage, precision, specificity, sensitivity, cost, throughput, quantification type, multi-matrix validation, evidence depth, and pQTL accuracy. Axes are identically ordered across panels so that platform profiles are directly comparable, and each platform’s characteristic strengths appear as the shape of its polygon. Scores are relative to the evaluated set and are justified with supporting citations in **Supplementary Tables S3** to **S9**.

#### Proteome Coverage, Precision, and Specificity

Scoring dimensions captured distinct and often opposing strengths across platforms. For example, proteome coverage ranged from focused panels of a few hundred analytes to broad assays reporting many thousands of proteins, with the Seer nanoparticle mass spectrometry and SomaSeq aptamer readouts providing the highest proteome coverage and the targeted NULISA proximity ligation panel providing the lowest. Precision and specificity frequently traded against coverage. For example, Somalogic’s aptamer assays have consistently shown the lowest reported coefficients of variation (median CVs of roughly 5% to 7% in independent cohorts), earning SomaSeq the highest precision score, but each aptamer recognises its target at a single binding site, raising concerns that single-site recognition is more vulnerable to off-target signal^19,20^. Head-to-head comparisons anchored to genetic and clinical evidence have generally favoured Olink’s dual-antibody design on target specificity^11,13^; however, in a split-sample comparison, Olink Explore HT showed a median CV of 35.7% against 6.8% for SomaScan 11k, with precision strongly dependent on whether analytes lay above the limit of detection^21^. Thus, while Olink scored highly on specificity, it scored the lowest of all platforms on precision (**Figure 2** and **Supplementary Table S3**).

#### Sensitivity and pQTL accuracy

The Alamar NULISA proximity ligation panel scored highest on sensitivity, reflecting its attomolar limits of detection^6^ and a widely replicated capability to measure ultra-low-abundance proteins, including phosphorylated tau species, in blood^22,23^. MS via Seer scored highest on pQTL accuracy because peptide-level identification can distinguish genuine abundance changes from the binding artifacts that inflate spurious associations on affinity platforms. Missense variants can still produce false signals on MS by producing peptides absent from the reference lib1rary; however, this can be accounted for by quantifying against peptide libraries that include variant alleles^15^.

#### Cost per sample

Nomic scored highest on cost per sample based on its publicized $50-per-sample price. Published pricing for the other platforms was sparse or absent, and variable where it existed. The remaining platforms were therefore scored on a combination of published evidence^24^ and direct quotes historically obtained by the authors. Alamar NULISA scored 4/5, with pricing generally in the low hundreds of dollars per sample, whereas Olink Explore HT, SomaSeq, Seer Proteograph XT, and Biognosys TrueDiscovery all scored 3/5, with pricing varying from the low-to-high hundreds depending on project size and scope.

#### Throughput

We scored throughput based on demonstrated scale in the published literature. Olink scored 5, having profiled 54,219 UK Biobank samples using its earlier Explore 3072 assay^2^, with a 600,000-sample second phase underway using Olink Explore HT. Nomic also scored 5, having profiled 7,392 samples on a 191-plex panel in under one week^7^. SomaSeq / Somalogic scored 4, as its largest published single study comprised 35,559 participants^25^, with larger deployments in progress but not yet published; this score may be revised based on future publications. Seer and Biognosys both scored 3, as MS acquisition limits throughput to tens of samples per day per instrument, and scaling requires instrument parallelization^24^. Alamar NULISA also scored 3. Although its workflow is fully automated^6^, published studies have yet to deploy the assay in more than a few thousand samples. Like SomaSeq, this score may be increased based on future published outputs of collaborations between Alamar and global biobanks (**Supplementary Table S6**).

#### Multi-matrix validation

The multi-matrix validation score reflects the breadth of sample matrices in which a platform has published validation, as well as how much re-validation a new matrix requires. Biognosys TrueDiscovery scored 5 on this metric because bottom-up DIA applies one workflow to any proteinaceous input that can be digested. Plasma, serum, fresh-frozen tissue, FFPE, and cell culture all use the same acquisition and detection chemistry^26,27,28,29,30^, and although measurements are still read out as relative concentrations, peptide sequence identification does not need re-validation in new matrices. By contrast, binder-based assays are validated primarily in biofluids, and antibodies or aptamers may face distinct specificity challenges in orthogonal matrices. Both SomaLogic and Olink have been widely deployed across CSF, tissues, cell lysates, urine, and more^5,31,32,33,34^, but the values they report are often not comparable between matrices. Somalogic readouts shift with sample type(s) and cross-specimen comparisons often require bridging^31,35,36^, and even paired serum and EDTA plasma have disagreed for 36 of 80 evaluable Olink proteins in a study of the 92-protein Olink Immuno-Oncology panel^37^; thus, both Somalogic and Olink scored 3. Similarly, Alamar NULISA is well-characterized in plasma and CSF^6,38^, but it is not validated in tissues, and therefore scored 3. Nomic nELISA scored 2 as it is validated mainly in cell culture and secretome studies to date^7^, while Seer, although MS-based, scored 3 as its nanoparticle corona chemistry is validated for biofluids only.

#### Quantification type

Quantification type was scored categorically. Platforms that deliver true absolute quantification in traceable physical units scored higher than those reporting relative or normalised signals, with none of the six platforms scoring 5 of 5. The Nomic nELISA assay scored 4 as it produces analyte concentrations in physical units (pg/mL), back-calculated from per-target recombinant standard curves run in the same multiplex format as the samples^7^; however, calibration is against recombinant antigen rather than certified reference material, so values are traceable to the manufacturer’s standards rather than to SI units and may diverge from endogenous proteoforms. All other affinity-based platforms scored 2 because their mid-to-high-plex offerings provide relative signal in arbitrary units normalised within analytes across samples and not convertible to physical concentration(s). The two MS-based platforms, Seer and Biognosys, scored 3. Although label-free MS intensity is relative, intensity-based estimators such as iBAQ^39^ and Top3^40^ can facilitate approximate within-sample ranking of protein abundance, and both platforms offer a documented route to true absolute quantification through isotopically labelled internal standards^41^ without a change of platform^42^.

#### Evidence Depth

Where the peer-reviewed literature permitted, scores were anchored to published measurements, with head-to-head comparisons weighted above single-platform characterisations; where it did not, we relied on manufacturer white papers and direct correspondence with vendors. The ‘evidence depth’ score indicates the degree with which a platform was evaluated based on peer-reviewed, fully quantitative evidence versus unreviewed or semi-quantitative evidence, thereby qualifying the confidence assigned to a platform’s remaining scores. Established platforms such as Olink and SomaSeq / Somalogic scored 5 due to their repeated characterisation by independent groups in large cohorts, such that the remainder of their scores rested on data we did not have to take on trust. Contrastingly, emerging platforms like Nomic nELISA scored lower, as most data were derived from unpublished analyses, manufacturer white papers, and direct correspondence with the vendor; thus, the platform’s remaining nine metrics should be treated as provisional. This ‘Evidence Depth’ dimension is not incorporated into any composite score, since confidence in a measurement is not an attribute of the quantity measured. Because publication volumes accrue with time on the market and installed base, the score indexes commercial maturity alongside technical merit, and would penalise newer platforms for little beyond their novelty if factored into an overall composite. Thus, readers reweighting the matrix for their own purposes should treat evidence depth as a modifier on the other nine scores and not as a tenth quantitative attribute.

### Protein-level coverage and platform-exclusive content

The availability of documentation for protein coverage varies between platforms. Some vendors provide their complete protein target databases via open web applications while others deliver their target lists upon request, often in exchange for marketing consent. To address these inconsistencies, we built a protein coverage layer that maps each platform’s content to UniProt identifiers and assembles a centralized, cross-platform catalogue facilitating streamlined protein lookups. Across the six platforms, this catalogue contained 30,147 assays targeting 15,123 protein identifiers. Most assays mapped to a single protein, although 15 Olink Explore HT assays each targeted two or more closely related proteins (such as MICA and MICB).; Additionally, certain single identifiers, such as tau (MAPT; P10636), could carry many assays including total tau (measured on five of the six platforms) and different forms of phosphorylated tau (measured on the NULISA Neuro220 panel).

The overlap structure across platforms was highly uneven (**Figure 3A**). Of the 15,123 protein identifiers, 6,720 (44.4%) were measured by only a single platform, and just 109 (0.7%) were shared by all six. This directly impacts study design, since a result observed on one platform may not be observable on another, and cross-cohort replication may be constrained by how little of the measured space the platforms share. Moreover, protein identifiers shared across platforms can still be assayed through different epitopes, producing discordant measurements of the same protein^11,12,13^, and further complicating independent replication.

**Figure 3.**
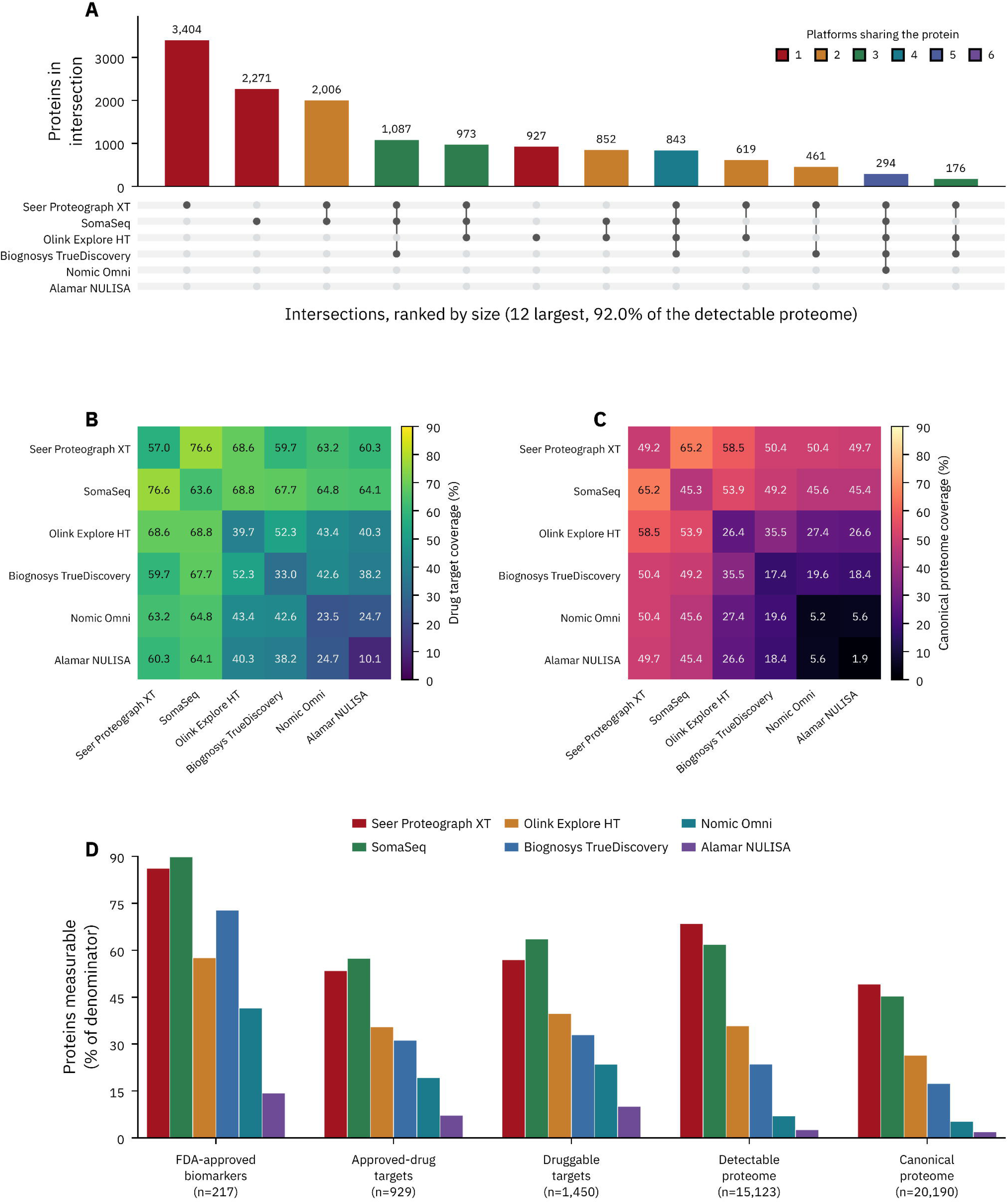
Proteome coverage across different denominators. Platforms are ordered throughout by descending catalogue size (Seer Proteograph XT, 10,364 identifiers, to Alamar NULISA, 387). (A) Overlap structure across the six platforms, shown as the twelve largest intersections ranked by size, which together account for 92.0% of the union. Of 15,123 distinct UniProt identifiers measurable by at least one platform, 6,720 (44.4%) are measured by a single platform and 109 (0.7%) by all six. Platform-exclusive content is concentrated in the discovery-scale platforms, with Seer Proteograph XT contributing 3,404 exclusive identifiers, SomaSeq 2,271 and Olink Explore HT 927, against 109 for Biognosys TrueDiscovery, 8 for Nomic Omni and 1 for Alamar NULISA. Bar colour denotes the number of platforms sharing each intersection. (B) Coverage of the 1,450 druggable targets and (C) of the 20,190-protein canonical proteome for each platform alone (*diagonal*) and for each platform pairing (*off-diagonal*). Colour scales are independent; both span 0 to 90%. Coverage here is defined as catalogue membership by UniProt identifier, and the two mass spectrometry catalogues are unions across studies, matrices and instrument configurations. (D) Proportion of five reference sets measurable by each platform alone, ordered from the most clinically stringent (217 FDA-approved protein biomarkers) through approved drug targets (929) and druggable targets (1,450) to the detectable proteome (15,123) and the canonical human genome-encoded proteome (20,190). The ordering between platforms inverts across denominators; for example, SomaSeq leads on the three target-based sets while Seer Proteograph XT leads on both proteome-scale sets.

Platform-exclusive contributions to the total assay count also ranged widely. Mass spectrometry via Seer Proteograph XT contributed 3,404 exclusive proteins (32.8% of its own catalogue) and the SomaSeq aptamer readout contributed 2,271 (24.3% of its own catalogue), followed by Olink Explore HT at 927 (17.1% of its own catalogue); thus, the three broadest platforms accounted for 98% of all single-platform content. The remaining three contributed far less exclusive content by count, including 109 exclusive proteins on Biognosys TrueDiscovery, eight on Nomic Omni, and one on Alamar NULISA (six when including phosphorylated tau species).

### Coverage of drug targets, clinical biomarkers, and the broader proteome

The UniProt database contains over 20,000 reviewed human entries - a collection informally referred to as the canonical human proteome^43,44^. However, this ‘canonical’ map understates the true complexity of the proteome by orders of magnitude, as sequence variation, alternative splicing, and post-translational modification expand it into hundreds of thousands to millions of distinct proteoforms^45,46,47^ distributed in blood plasma across a concentration exceeding ten orders of magnitude^48^. The six platforms assessed in this study collectively captured 15,123 proteins, representing just under 75% of the canonical human proteome and a small fraction of the broader proteoform landscape. Of these, approximately 1,450 are targets of approved or clinical-stage drugs ^49^, including 929 targeted by at least one approved drug, while a separate set of 217 proteins are measured by FDA-cleared or approved clinical assays as diagnostic biomarkers^50^.

SomaSeq and Seer both captured the most substantial fractions of drug targets (63.6% and 57.0% respectively; 76.6% combined) (**Figure 3B**). However, of the 1,450 proteins targeted by approved or clinical-stage drugs (hereafter referred to as the “druggable proteome”), 293 (20.2%) were not detected by any of the six platforms, and a further 30.6% were detected by only one measurement class (MS or affinity proteomics), indicating that combining MS with one or more binder assays may be necessary to adequately power proteomic drug target validation studies. As an illustration of modality-specific coverage, targets of approved and clinical-stage antineoplastic kinase inhibitors (89 target genes) and monoclonal antibodies (50 target genes) were well covered, with the six-platform union capturing 97.8% and 92.0% respectively, led by SomaSeq at 92.1% of kinase inhibitor targets. Conversely, all six platforms combined detected only 30.8% of 156 druggable ion channels (the family targeted by common anaesthetics and antiarrhythmics) and 47.4% of 154 G-protein-coupled receptors (the family targeted by beta-blockers, antihistamines, and most antipsychotics) (**Figure 6A**). Affinity panels lack sufficient reagent coverage for most of these multipass membrane protein families, while MS detects these families only at low endogenous plasma abundance, likely underpinning their low coverage rates. Overall, affinity panels captured protein classes that their antibodies or aptamers were primarily designed to detect - measuring 94.8% of 192 membrane receptors versus only 66.7% detected by MS. The same pattern held for secreted ligands (89.2% affinity against 68.4% MS across 158 proteins) and nuclear receptors (91.7% against 33.3% across 24). MS showed much stronger coverage of transporter proteins, capturing 75.6% of all transporters against 43.3% for affinity panels (**Supplementary Figure S4** and **Table S14**).

The ‘detectable’ proteome, representing the union of 15,123 proteins measured by at least one of the six platforms, was well captured by SomaSeq and Seer at 61.8% and 68.5%, respectively, and 92.9% in combination. Separately, these two platforms captured smaller segments of the canonical human proteome (45.0% for SomaSeq; 48.9% for Seer), indicating that combining the two may be necessary to maximize coverage. However, at 64.7% combined coverage, this would still leave over one third of the canonical proteome unmeasured; adding a third high-plex platform such as Olink Explore HT would raise coverage only to ∼69.5%, highlighting opportunities to close the measurement gap as part of future assay expansions (**Figure 3C**).

All six platforms covered FDA-approved clinical biomarkers well for their size, but the smallest panels were the most enriched. 214 of the 217 FDA-approved biomarkers are measurable by at least one platform, so a random panel’s expected count is its size times 214/15,123. Relative to a random draw of equivalent size from the detectable proteome, Nomic Omni 1000 was enriched six-fold (90 FDA biomarkers observed against 15 expected) while Alamar NULISA’s combined Neuro220 and Infl250 panels were enriched 5.7-fold, against 1.5-fold for SomaSeq and 1.3-fold for Seer Proteograph XT (all *p*<1×10^−10^, hypergeometric) (**Figure 3D**). Absolute coverage rose with panel size (Spearman rho = +0.89) while enrichment fell (rho = −0.94); as we expected, since a larger panel covers more of the fixed 217 but dilutes its biomarker fraction. Nomic Omni measured 1,058 proteins, roughly a tenth of SomaSeq’s content, yet measured 41.5% of the FDA-approved biomarker set against SomaSeq’s 89.9%, and 8.5% of everything Nomic measured was an approved biomarker - four times SomaSeq’s 2.1%. This enrichment strongly reflects panel design, since focused panels like Nomic’s and Alamar’s are assembled around established clinical analytes and candidate biomarkers. This reflects an advantage over broader platforms like Seer, SomaSeq, and Biognosys only when users explicitly prioritize cost, absolute quantification, or sensitivity. Otherwise, the broadest panels offer the highest coverage of clinical biomarkers.

### Help Me Choose: A proteomics technology recommendation tool

Help Me Choose is a dedicated technology recommendation tool guided by the atlas’s nine quantitative scoring dimensions and the relative weights an investigator assigns to those scores in real-time. A researcher specifies a study context (primary research goal, sample matrix, and study size), sets seven priority sliders (cost, proteome coverage, precision, specificity, sensitivity, throughput, and pQTL accuracy), and answers two required capability questions (central nervous system (CNS) focus and/or longitudinal design) with two further optional ones (absolute quantification and/or post-translational-modification detection) (**Figure 1A**). The tool converts these inputs into an effective weight for each dimension, applies eligibility filters, and returns a ranked, annotated list of platforms based on these user preferences and the available evidence base (**Figure 1B**). The effective weight calculation, two-pass calibrated scoring, and hard filters are described in full in **STAR Methods**.

We tested whether the tool structurally advantages any platform by enumerating every allowed combination of user inputs. This grid included six goals, six sample matrices, four study sizes, four capability states – every combination of the two binary study characteristics (CNS/pTau biomarker focus and longitudinal/repeat-sampling design), each yes or no – and three weight levels for each of the seven sliders. This produced 6 × 6 × 4 × 4 × 3^7^ = 1,259,712 scenarios, each evaluated deterministically. Each combination was weighted by its real-world prevalence, estimated from the inputs of 65 research scientists (Figure 4B; STAR Methods). In each scenario, we recorded the top-ranked (“winning”) platform. Because a platform can either win outright (**Figure 4A**) or share a tie (**Figure 4C**), we recorded both statistics. No single platform was recommended outright in 62.9% of simulated scenarios, closely matching the 65.1% of real-world sessions (41 of 63) that ended in a co-recommended tie. Because most scenarios end in ties, we report the co-recommendation rate as the primary summary of platform performance, with outright wins reported as a secondary result.

**Figure 4.**
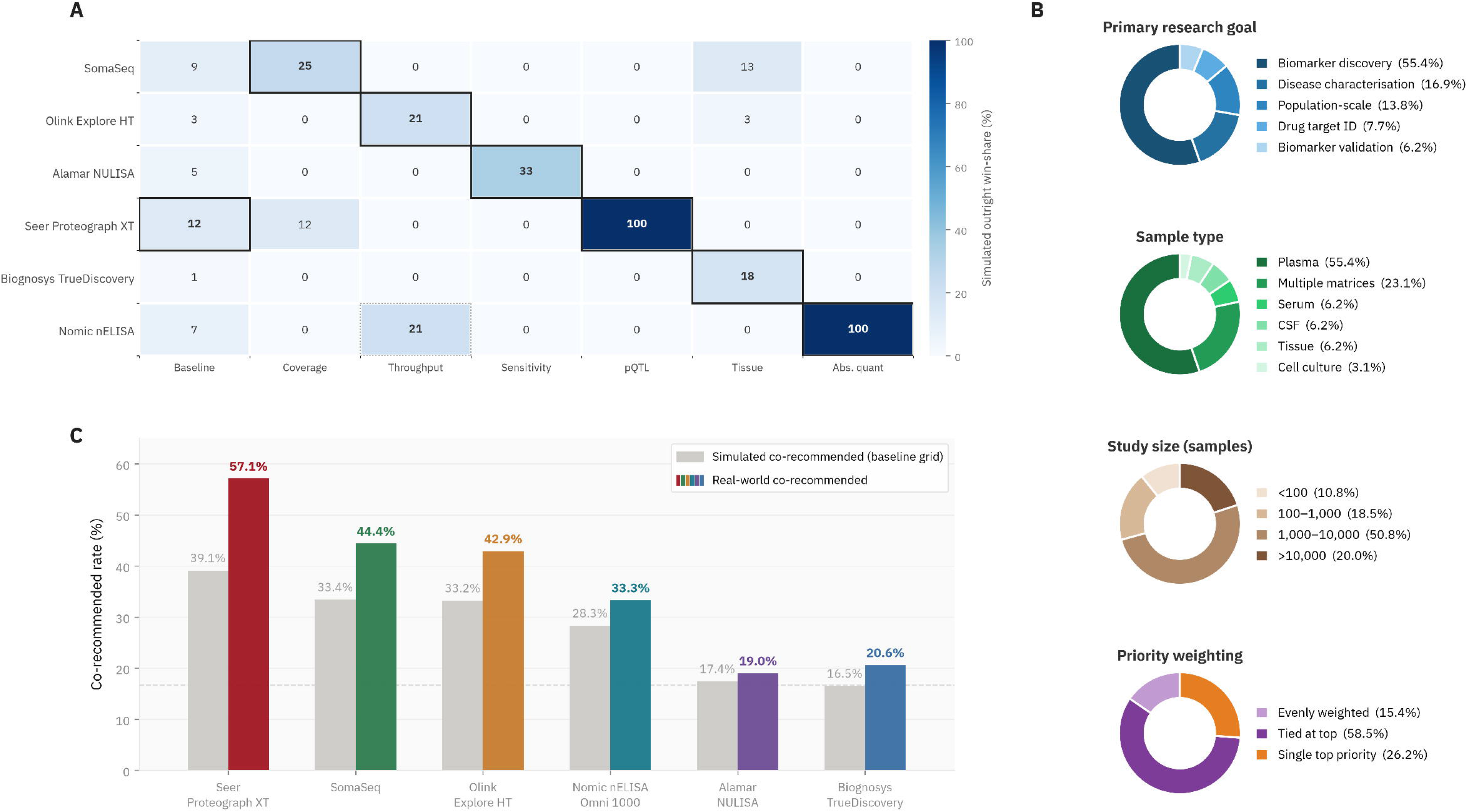
Help Me Choose: fairness, real-world demand, and validation. (A) Platform-by-study-context matrix of simulated outright win-share (%) from the 1,259,712-scenario fairness sweep; each platform leads where its strengths are decisive; for example, Biognosys on tissue, NULISA on sensitivity, Nomic on absolute quantification. (B) Profile of recorded researcher requests (primary research goal, sample type, study size, and priority weighting), the real-world demand that grounds the fairness analysis and informed tool calibration. (C) Simulated versus real-world co-recommended rate per platform, based on 63 scored study profiles. The analogous outright-win comparison is reported in **Supplementary Figure S5**. Displayed separations are small and most profiles resolve to ties, with no single platform dominating.

Seer Proteograph XT led on both measures in the baseline simulated scenario (39.1% co-recommended, 12.3% outright), followed by SomaSeq (33.4%, 9.4%), Olink Explore HT (33.2%, 3.1%), Nomic Omni (28.3%, 6.7%), Alamar NULISA (17.4%, 4.7%) and Biognosys TrueDiscovery (16.5%, 1.0%). We then maximised one priority slider at a time, holding the others at baseline and restricting to plasma so sample type could not affect the result. Each dimension produced a different winner, reported as the share of scenarios in that slice won outright: SomaSeq for coverage (25.0%), Alamar NULISA for sensitivity (33.3%), and Seer Proteograph XT for pQTL accuracy (100.0%, the only dimension with a unanimous winner). Throughput ended in a tie between Olink Explore HT and Nomic Omni (20.8% each), while cost favoured Nomic Omni 1000 outright (**Figure 4A**). Conditioning on study context showed Biognosys TrueDiscovery and Alamar NULISA, the two platforms with the lowest baseline share, leading where their strengths are decisive: Biognosys in tissue matrix studies (17.8% outright win rate when restricted to tissue) and NULISA in neuroscience- and CSF-focused studies, alongside Nomic in absolute-quantification studies (**Figure 4A**).

Real-world respondents most commonly designed their proteomics studies around plasma matrices (55.4%) with a primary research goal of biomarker discovery (55.4%) in more than 1,000 samples (70.8%) (**Figure 4B**). As a secondary fairness check, we restricted the grid to this modal real-world profile. With goal, matrix, and capability questions fixed, and study size restricted to the two levels above 1,000 samples, only the seven slider weights (three levels each) varied, giving 2 × 3⁷ = 4,374 scenarios. In this more restricted set of simulations, Seer Proteograph XT led (45.0% co-recommended, 17.6% outright), ahead of Olink Explore HT (39.7%, 4.1%), Nomic Omni (36.7%, 9.3%) and SomaSeq (29.1%, 6.7%). Alamar NULISA (8.1%, 0.1%) and Biognosys TrueDiscovery (0.0%, 0.0%) trailed; Biognosys because its tissue matrix strength is irrelevant in a plasma-only context, and NULISA because it provides focused panel coverage.

Our “Help Me Choose” tool sets any absolute quantification requirement as a hard filter. Among the six scored platforms, only Nomic nELISA reports validated physical-concentration (pg/mL) output, so it won in 100% of scenarios where absolute quantification was required. In observed survey usage, this requirement was specified as “required” in 10.8% of sessions, “preferred” in 50.8%, and “not needed” in 38.5%. Where Nomic is recommended, the tool informs users that several excluded platforms offer absolute quantification through separate, targeted products not included in our analysis (e.g., Olink Flex/Target 48, Alamar NULISAseq AQ, or Biognosys TrueSignature).

Re-scoring the 63 eligible real-world sessions through the production app (STAR Methods), Seer Proteograph XT led (57.1% co-recommended, 15.9% outright), ahead of SomaSeq (44.4%, 0.0%), Olink Explore HT (42.9%, 0.0%), Nomic Omni (33.3%, 19.0%), Biognosys TrueDiscovery (20.6%, 0.0%) and Alamar NULISA (19.0%, 0.0%) (**Figure 4C**). Four platforms recorded no outright wins, consistent with the rarity of outright wins overall (34.9% of sessions).

### Help Me Combine: A tool to help rank proteomics platform pairings

#### Tool overview

Help Me Combine is a combination recommendation tool designed for investigators who intend to deploy two proteomics platforms and wish to determine which pairing adds most to their study. The tool operates over all six assays and therefore over fifteen pairs, scoring each pair on seven axes (coverage, complementarity, combined measurement score, cost, class balance, sample volume, and PTM or proteoform detection capability). On each axis, the fifteen pairs are rescaled to a 0-to-1 range, with the lowest-scoring pair at 0 and the highest at 1. A pair’s overall score is then the weighted average of its rescaled axis values.Users select up to three of eleven priorities. Five are axis priorities (coverage, complementarity, low cost, low sample volume, and tissue-specific study design) that set their axis weight to the maximum; six are measurement priorities (precision, specificity, sensitivity, absolute quantification, pQTL accuracy, and throughput) that raise the combined measurement score axis and restrict that score to the selected properties. The full specifications for this tool are provided in STAR Methods (Platform combination tool) and outlined in **Figure 5**.

**Figure 5.**
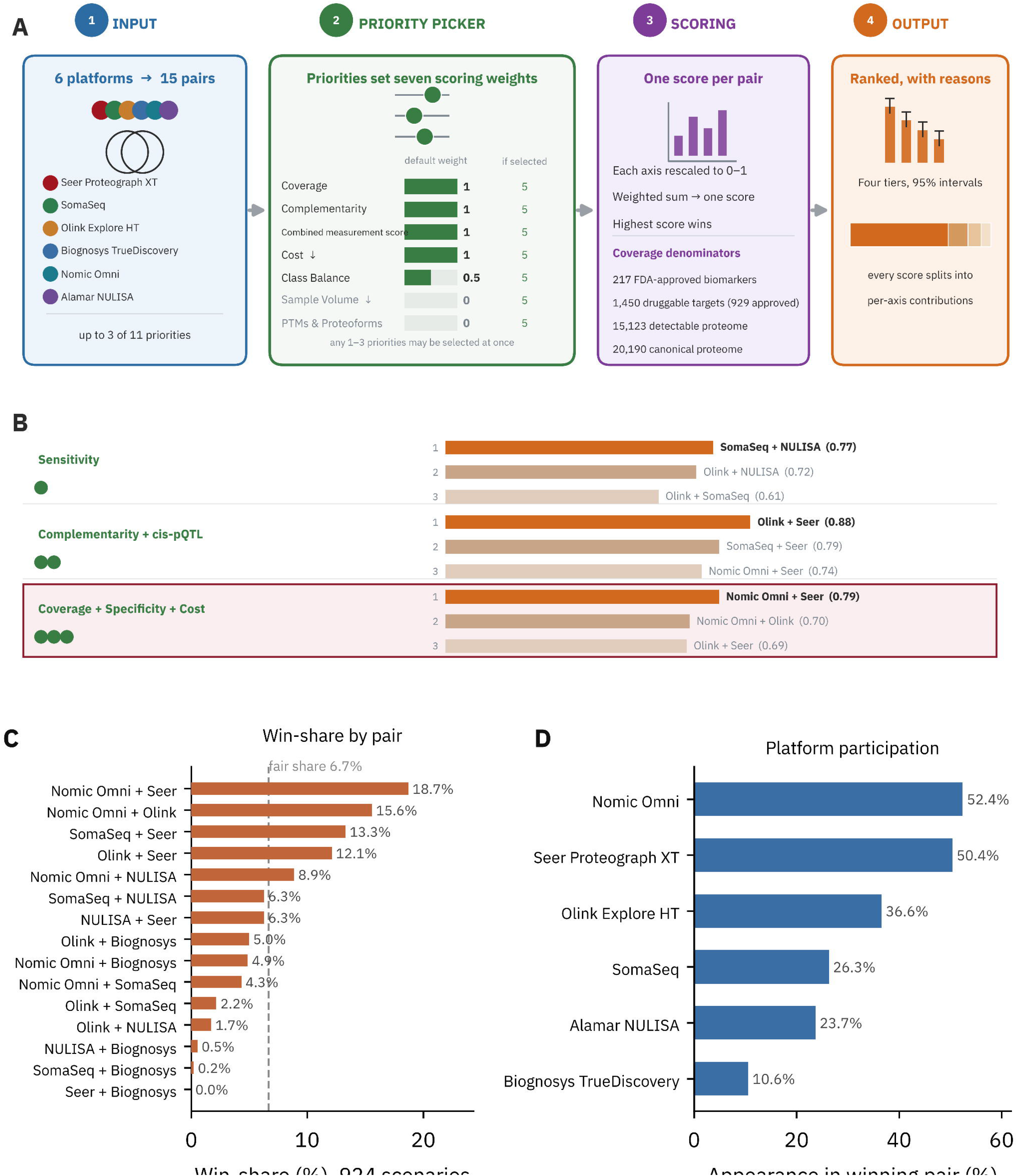
Structure and behaviour of the Help Me Combine recommendation tool. (A) The tool in four stages. Six platforms yield 15 candidate pairs. Users select up to three of eleven priorities, each of which raises one of seven scoring axes from its default weight to 5 (coverage, complementarity, combined measurement score, cost, class balance, sample volume, and PTM and proteoform capability. Cost and sample volume are inverted so that a lower value scores better, and sample volume and PTM and proteoform capability are held at zero weight unless selected). Combined measurement score represents the average of the two platforms on each selected property, meaning a pair scores well only if both platforms are individually strong. With no combined measurement priority selected, that average runs over five intrinsic properties (precision, specificity, sensitivity, quantification type, and cis-pQTL accuracy); three further properties reflecting platform maturity (throughput, evidence depth, and multi-matrix validation) sit at zero weight until a user selects them. On each axis, the fifteen pairs are rescaled to a 0-to-1 range, with the lowest-scoring pair at 0 and the highest at 1. A pair’s overall score is then the weighted average of its rescaled axis values. Coverage is reported against four numeric denominators (1,450 ChEMBL drug targets, of which 929 are targets of approved drugs; 217 FDA-approved biomarkers; 15,123 proteins measurable by at least one platform in this atlas; 20,190 canonical human proteins) and a qualitative PTM and proteoform capability flag. (B) Tool output for five priority selections against the druggable target denominator, showing the three highest-scoring pairs and their calibrated scores. With no priority selected, the leading pair is Olink Explore HT with Seer Proteograph XT; selecting coverage returns SomaSeq with Seer Proteograph XT, cost returns Nomic Omni with Olink Explore HT, and coverage with precision together return SomaSeq with Alamar NULISA. The highlighted row shows an example of the best combination for three key priorities (coverage, specificity and cost), won by Nomic Omni with Seer Proteograph XT, which wins none of those three priorities individually - ranking fourth, second, and third on them respectively.

#### Complementarity score and combined drug target coverage

The Help Me Combine tool defines complementarity as the net-new content a second platform adds to the first, derived from the protein coverage lists described previously. In a scenario where a user weights coverage highly, the leading combination is SomaSeq with Seer Proteograph XT, reaching 76.6% of the druggable target list (**Figure 6B**) and adding 188 net-new targets (13.0%) beyond the stronger single platform. However, two platform pairs with very similar coverage can still notably differ on the second platform’s added content. For example, Nomic Omni with SomaSeq detects 64.8% of druggable targets but adds only 17 net-new proteins, because the smaller panel is largely a subset of the larger; Nomic Omni with Seer Proteograph XT detects slightly less at 63.2% but adds 91 proteins from an orthogonal technology. Coverage and complementarity therefore rank these two pairs in opposite orders.

**Figure 6.**
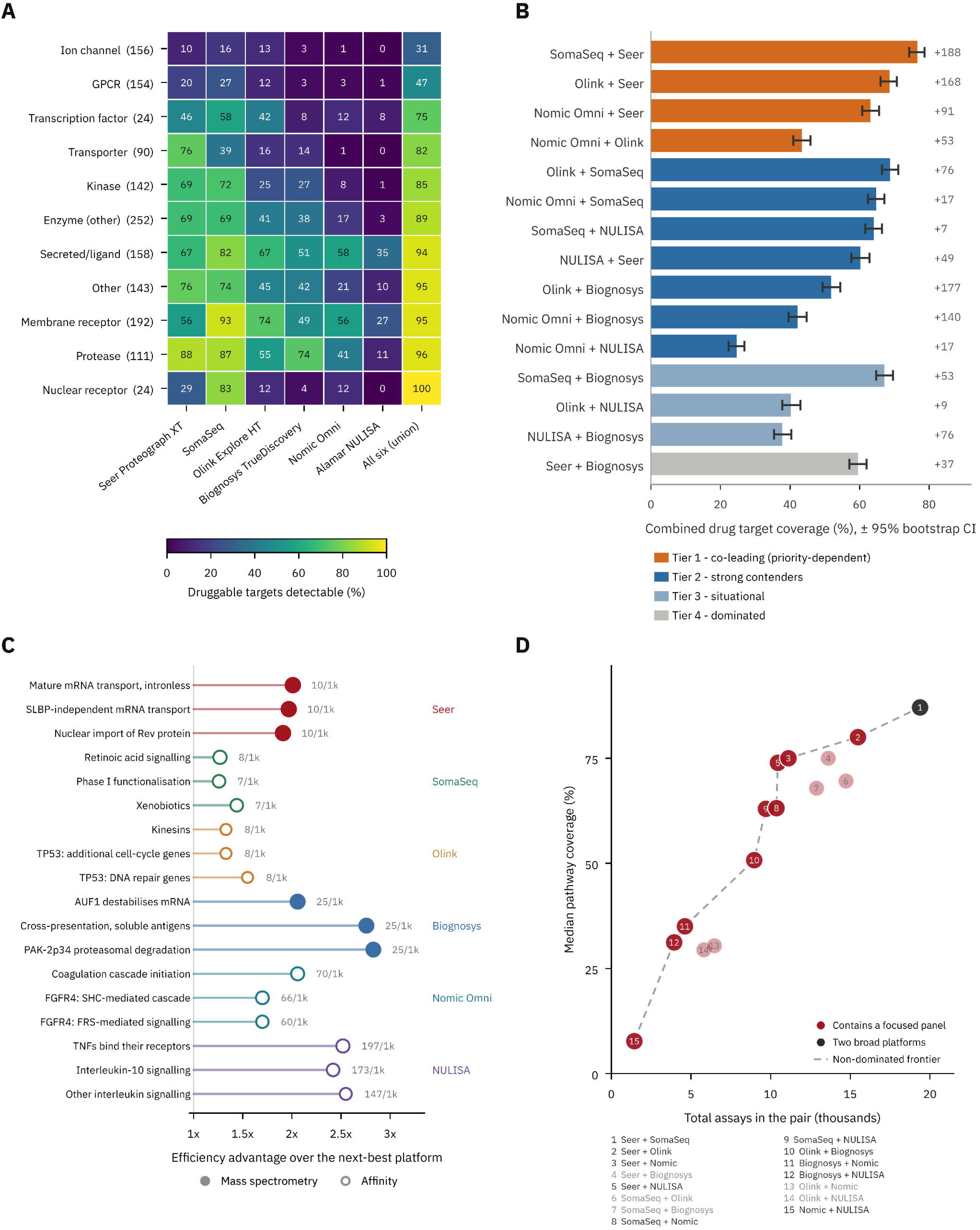
Coverage of drug targets and biological pathways, by platform and by pair. Top row, druggable targets; bottom row, Reactome pathways; in each row, single platforms first, then pairs. (A) Percentage of each druggable target class detectable by each platform, for the eleven classes and 1,450 targets of the druggable set; class sizes are given in parentheses. Rows are ordered by union coverage across all six platforms, so the least accessible classes read first. Ion channels (best single-platform coverage 16%) and G-protein-coupled receptors (27%) are the least covered, membrane receptors (93%) and proteases (88%) the best; the per-class leader alternates between SomaSeq (seven classes) and Seer Proteograph XT (four). (B) Combined druggable target coverage for all 15 pairs, ordered by robustness tier and then by coverage within tier, so the strongest pair reads at the top; error bars are 95% bootstrap confidence intervals. Tier 1 (orange) are the four pairs highest by priority-sweep win share and are treated as co-leading. Tier 2 (blue) are seven further pairs that are not Pareto-dominated. Tier 3 (light blue) are three pairs that win no scenario but remain non-dominated. Tier 4 (grey) is the single Pareto-dominated pair, matched or exceeded by another pair on every scored axis. Right-hand labels show net-new druggable targets added by the second platform beyond the higher-coverage member. (C) The three Reactome pathways each platform covers most efficiently, plotted as its coverage per 1,000 assays divided by that of the next-best platform on the same pathway. A value of 2x means the platform reaches that pathway twice as efficiently as any competitor. Absolute efficiency is printed beside each marker. (D) Total assays against median pathway coverage for all 15 pairs, with the non-dominated frontier joined by a dashed line. Points are numbered by descending coverage and named in the key below; greyed key entries signify dominated pairs. Nine of the ten frontier pairs contain a focused panel.

#### Combined pathway coverage

Of 1,285 Reactome pathways testable against the canonical proteome, 613 carried at least twenty members measurable by at least one platform. Absolute coverage of these pathways was dominated by the two largest catalogues, SomaSeq and Seer Proteograph XT, which together led 612 of the 613 pathways. That ranking, however, follows directly from panel size; it says nothing about whether smaller panels are enriched for particular biological pathways, nor does it help users choose an efficient platform combination under a constrained budget. To assess this, we normalised by panel size, expressing coverage as percentage points of pathway coverage per 1,000 assays. On this measure, the most efficient platform was a focused panel for 489 of the 613 pathways. Alamar NULISA reached 197 points per 1,000 assays on TNF receptor binding, against a best competing value of 78, and led interleukin-10 and other interleukin signalling; Biognosys TrueDiscovery led proteasomal and mRNA-destabilisation pathways at 2.1 to 2.8 times the next-best platform (**Figure 6C**). Pairing a focused panel with a broad platform kept this efficiency on pathway coverage while expanding proteome coverage. We plotted every platform pair by its total assay count against the median coverage it achieved across the 613 pathways (**Figure 6D**). Ten of the fifteen pairs offered a combination that no other pair beat on both counts at once (more coverage for fewer assays) and nine of those ten included a focused panel.

#### Fairness testing and uncertainty reporting

We tested whether the pairing tool structurally favours any pair or platform using a deterministic priority sweep that enumerated all 231 combinations of one to three priorities across the four coverage denominators, comprising 924 scenarios in total (**Supplementary Figure S6**; STAR Methods). No pairing recommendation exceeded 18.7% of scenarios. Nomic Omni with Seer Proteograph XT led (18.7% of scenarios), followed by Nomic Omni with Olink Explore HT (15.6%), SomaSeq with Seer Proteograph XT (13.3%), and Olink Explore HT with Seer Proteograph XT (12.1%), with a tail to 0.0% (Supplementary Figure S6A). Fourteen of the fifteen pairs won at least one scenario (all but Seer Proteograph XT with Biognosys TrueDiscovery). Every platform appeared in a substantial share of winning pairs (Nomic Omni 52.4%, Seer Proteograph XT 50.4%, Olink Explore HT 36.6%, SomaSeq 26.3%, Alamar NULISA 23.7%, Biognosys TrueDiscovery 10.6%) (**Supplementary Figure S6B**).

Nomic Omni and Seer Proteograph XT each appeared in roughly half of winning pairs; Nomic Omni for its cost, throughput, and absolute quantitation, and Seer Proteograph XT for its broad coverage and pQTL accuracy. Alamar NULISA (23.7%) was recommended mainly where sensitivity was prioritised. Biognosys TrueDiscovery appeared least (10.6%): its multi-matrix flexibility was decisive only when tissue work was prioritised, and 87.1% of its published catalogue overlaps with Seer Proteograph XT, so it rarely added coverage a broader pair could not. Unlike the Help Me Choose tool’s prevalence-weighted baseline, the priority sweep weights every combination equally, since no real-world priority selections have been collected since the tool’s launch in July 2026.

### An open, reproducible web resource

APT is a free web application available at https://aptatlas.org, including an interactive platform comparison section, both decision tools, a protein coverage browser enabling lookups of specific proteins or target lists, and the curated evidence database that links each score to the head-to-head and characterisation studies supporting it. The resource is a browser-based application (built with the React JavaScript framework) compiled to a self-contained HTML file. Both of its recommendation tools execute in the browser, so the scoring itself does not depend on a server. Study context and priority selections are recorded, with consent, to characterise real-world demand. The analysis in this Resource uses those responses in aggregate. Platform metadata and scoring data are held as version-controlled configuration files separate from the application logic, which allows scores to be updated as evidence appears and makes each release a citable state of the resource. The version described here is v4.0 (7 September 2026).

## Discussion

Research into plasma and serum proteomics in cohort and biobank settings has grown approximately tenfold in a decade, from 66 PubMed-indexed papers in 2015 to 692 in 2025. Protein QTL studies grew from 4 papers to 346 over the same period, underscoring an exponentially expanding base of human geneticists and molecular epidemiologists increasingly relying on proteomics for large-scale discovery and translation. This growth has been accompanied by proliferation of viable platforms. At least six mid-to-high-plex proteomics services are now commercially available, spanning affinity and mass spec families and a 26-fold range in panel size, from 385 assays to more than 10,000 proteins. Multiple head-to-head benchmarking studies have evaluated the strengths and weaknesses of these technologies, but each has compared a specific set of platforms on one cohort, matrix or endpoint, requiring investigators to reconcile evidence from divergent sources and extrapolate to a study design that no source directly addresses. In the absence of a consolidated comparison, platform choice is frequently determined by precedent, investigator expertise, or existing vendor relationship(s). These determinants are rarely revisited as products evolve, and they leave no citable record of why a cohort was profiled with one technology versus another. APT addresses this gap with a versioned, curated synthesis of the published evidence in which the weighting behind every platform recommendation is explicit and adjustable. The resource is engineered to provide fair, transparent guidance on which platform, or platforms, to deploy across 576 distinct study settings – from small longitudinal CSF studies searching for novel neuroscience biomarkers to plasma-based drug discovery studies in upwards of 10,000 participants.

Applied across those settings, the framework shows that no platform or pair is uniformly best. The leading choice shifts with the reference set used to judge coverage, and shifts further with the priorities an investigator specifies. SomaSeq offers the best coverage of drug targets, whereas Seer Proteograph XT offers the strongest coverage of the canonical human proteome (48.9% against 45%). Coverage of FDA-approved biomarkers is high for most platforms relative to their overall coverage, reflecting how most approved biomarkers are abundant, secreted plasma proteins^51^; even the smallest panels are strongly enriched for these markers. However, coverage of the broader proteoform landscape, comprising hundreds of thousands to potentially millions of distinct protein species^45,47^, remains out of reach for all current platforms. Thus, the same coverage number can be impressive or negligible depending on what it is measured against; most of the druggable proteome, a fraction of the canonical proteome, or almost none of the full proteoform space. “This platform measures *N* proteins” only becomes meaningful when *N* is stated as a share of the proteins that matter for the study at hand.

Our analysis revealed that a substantial proportion (20.2%) of the druggable proteome is undetected by any platform evaluated, and a further 30.6% is currently measurable only by MS or an affinity platform but not both, indicating that the two measurement families are non-redundant, and a combined approach should strongly be considered for drug discovery and target validation contexts. The fraction of the proteome undetected by existing technologies is highly concentrated in two multipass membrane protein classes, including ion channels (only 30.8% of which are detected by the six platforms) and G-protein-coupled receptors (only 47.4% of which are detected by existing platforms, despite representing over one third of all approved drugs^52^). These gaps reflect the limits of current assay chemistry^53,54,55^ and highlight areas where new reagents could substantially enhance proteogenomics-guided therapeutic target discovery and drug development^56^.

Proteome coverage represents only one of ten scoring dimensions, and in many contexts, may not reflect the decisive factor. For example, an investigator focused on reliable detection of brain-derived p-tau217 alongside other candidate CNS biomarkers at scale may prioritize the sensitivity of the Alamar NULISA assay over the breadth of Seer Proteograph XT. A researcher building population reference curves for a series of established and emerging clinical biomarkers may select Nomic Omni over Olink Explore HT, giving up the breadth of the latter for the calibrated quantitation capabilities of the former. For deep, multi-matrix work spanning both fluids and tissues, Biognosys TrueDiscovery may be preferred over SomaSeq, since untargeted DIA measures whatever proteins are present in a given matrix, including tissue-restricted content absent from fixed panels, and can enable post-hoc detection of PTMs that most affinity assays do not resolve. None of these scenarios favour the broadest catalogue. Each prioritises a different dimension of APT’s scoring framework, and each dimension is weighted according to the investigator’s stated preferences.

No platform or pair prevailed in more than a quarter of the contexts we evaluated, and the leading choice changed with the requirements of each study. The most frequently recommended single platform led outright in 12.3% of over 1.2 million exhaustive permutations, and the highest-ranked pair in 18.7% of 924 priority scenarios. These statistics show that the tools do not structurally favour any one platform and could, in that narrow sense, be considered fair. However, they cannot show that a given recommendation is correct; each platform wins where the dimensions it scores highest on are the ones a user prioritises, but this does not indicate that following the recommendation produces a better study. A benchmarking study evaluating all six platforms in the same cohort, comparing what each choice of individual platform and platform pairing yields in terms of novel biomarkers or aetiological candidates, would provide more robust data against which the APT scoring framework could be comprehensively validated.

Although both APT recommendation tools were designed to provide unbiased recommendations, real-world respondents received co-recommendations for Seer Proteograph XT, SomaSeq, and Olink Explore HT 10 to 18 percentage points more often than our fairness simulations projected. This likely reflects the composition of the respondent pool; stated study designs skewed heavily toward large or very large cohorts (70.8% of cases) and biomarker discovery or disease characterization goals (72.3%), which reward coverage and throughput strengths, while the tissue- and neuroscience-specific contexts that drove TrueDiscovery’s and NULISA’s simulated advantages were less common (tissue 6.2% and CNS focus 7.7% of sessions).

Several methodological considerations should guide how APT is used and how its scores are interpreted. Scores are ordinal 1 to 5 judgements, anchored to published measurements where the literature permitted and to vendor documentation and expert assessment where it did not; they are not continuous measurements, and platforms were scored at 3/5 across several metrics, representing par for the field. No score is fixed, and each will be regularly revisited as new evidence emerges. Coverage was defined using catalogue membership by UniProt identifier, which may be prone to over-counting usable coverage, since a nominal assay counts as covered even if it underperforms or falls below platform-specific limits of detection. The two MS platforms are study- and library-dependent, so their total protein counts are not strictly comparable to the fixed-content affinity panels. Notably, the Biognosys TrueDiscovery list was sourced from a published depletion workflow study and is a conservative 3,575-protein lower bound^18^; a protein corona-based “P2” enrichment workflow can increase coverage to approximately 7,000 proteins, but this list was not accessible at the time of analysis. SOMAmer platform scores were anchored to the published evidence for the array-based SomaScan platform rather than the newer SomaSeq readout, whose validation is still emerging, and should be revisited as that evidence accrues. Reactome pathway membership was denser for extensively studied biology across all platforms, likely reflecting an ascertainment bias arising from vendors choosing to design platforms that cover pathways of interest to customers; thus, a platform could lead on a sparsely annotated pathway even if it does not measure that pathway’s biology comprehensively. Finally, harder-to-quantify qualities, such as how readily a discovery finding converts to a custom targeted panel, are not yet scored, but may be integrated in future updates.

Proteomic platform selection is a methodological decision on the order of choosing between, or combining, genotyping arrays, sequencing panels, and whole-genome assays. It determines the breadth, depth, and comparability of the biological information a study can generate. However, unlike their genomic counterparts, proteomic data are less readily imputed to common reference panels or easily harmonized post-hoc. Affinity platforms agree only modestly at the level of individual proteins, and less than one percent of proteins are commonly detected across all six commercial platforms. Technology selection therefore commits investigators to a particular segment of the proteome, recorded in units that do not readily translate to other established or emerging platforms. Thus, platform choice warrants the same care given to genomic platform selection, and should be reported with the same specificity by recording the reference set, the analytical priorities, and the version of the evidence behind it. We built APT to enable straightforward production and scrutinization of this selection record. Because the underlying evidence changes faster than any static comparison can accommodate, platform scores are held as versioned configuration, reviewed monthly against new evidence, and released as citable snapshots. The evidence base is expanding at a sufficiently rapid pace that the framework should sharpen with each release, and platform choice should become progressively better informed.

## Acknowledgements

We are deeply grateful to the 65 anonymous scientists who answered the early version of the “Help Me Combine” tool, providing us with valuable real-world evidence to help refine and iterate the tool. We thank the proteomics platform vendors who provided protein target lists and pricing information. No vendor had any role in the design of the scoring framework, the assignment of scores, or the decision to publish. C.D.W. is thankful to Laura Winchester, Laura Ibanez, Marta Del Campo, Samantha Hutten, Zimple Kurlawala, Harris Bell Temin, Tyler Fortuna, Josh Denny, Geoff Ginsberg, Benoit Lehallier, and Martijn Kolijn for early feedback on the tool. We are indebted to the authors of the head-to-head platform comparison papers for providing a robust evidence base on which this tool could be built.

## Author contributions

Conceptualization, C.D.W.; methodology, C.D.W. and K.S.-B.; software, C.D.W.; formal analysis, C.D.W.; investigation, C.D.W.; data curation, C.D.W. and K.S.-B.; writing - original draft, C.D.W.; writing - review & editing, C.D.W. and K.S.-B.; visualization, C.D.W. Both authors critically reviewed the manuscript and take responsibility for the methods, scores, and conclusions.

## STAR Methods

### Key resources table

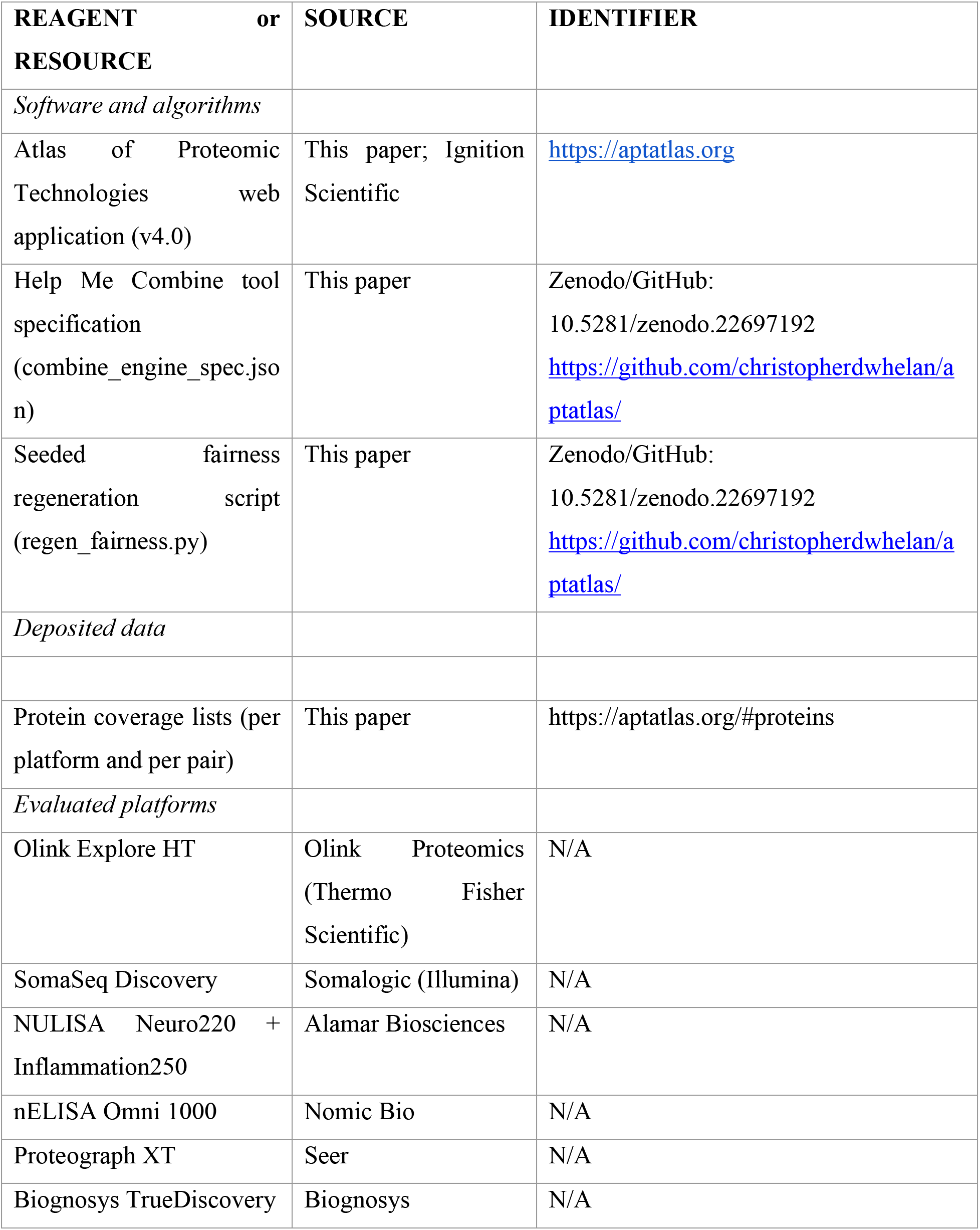

### Resource availability

#### Lead contact

Further information and requests for resources should be directed to the corresponding author, Christopher D. Whelan

#### Materials availability

This study did not generate new unique reagents; it evaluates commercially available proteomics platforms and does not involve materials that can be requested.

#### Data and code availability

The Atlas of Proteomic Technologies is available as a free, interactive web application at https://aptatlas.org (version 4.0, 7 September 2026).

The analyses reported here can be rerun from a downloadable bundle available on the app’s “Reproducibility” tab, which contains the tool specifications, the scripts that regenerate every fairness statistic, and the sweep results. The platform combination bundle contains the exact tool specification (combine_engine_spec.json, including the platform scores, per-platform and per-pair coverage values, axis definitions, and denominators), the regeneration script (regen_fairness.py, a self-contained replica of the production tool), and the fairness sweep result file (fairness_priority_sweep.csv). The sweep is a complete enumeration of every priority combination, so any researcher running it obtains the same result. The protein coverage lists that define platform and pair coverage against the druggable target, detectable proteome union, and canonical proteome denominators are provided in the same location. A third bundle provides the pathway analysis behind **Figure 6C**, **6D** and **Supplementary Table S10**, detailing the Reactome pathway-to-gene mapping, per-platform enrichment across all 1,285 testable pathways (reactome_enrichment_by_platform.csv), per-pathway coverage for all fifteen pairs (pathway_pair_coverage_full.csv), the two panel source files, and the class-level coverage table. Pathway coverage there is computed against the canonical proteome rather than the detectable proteome union, since restricting the background to measured proteins would make every pathway member detectable by construction.

Both fairness sweeps are complete enumerations, ensuring any investigator that runs them obtains the same results. The Help Me Combine tool’s scoring algorithm is stored in a self-contained JSON file, so its analysis can be rerun without the application source. The Help Me Choose tool’s weighting logic is kept in the application itself, so reproducing its sweep requires the released source.

The tool specification and seeded scripts have been deposited on Zenodo and mirrored on GitHub under a citable DOI: 10.5281/zenodo.22697192. Any additional information required to reproduce the analyses reported here is available from the lead contact on request.

### Method details

#### Resource development

The APT scoring framework and recommendation tools were developed iteratively, combining published evidence, expert judgement, and real-world researcher feedback (**Figure 1A**). We first curated a base of head-to-head platform comparison studies including investigations in plasma, serum, cerebrospinal fluid, and other sample matrices. A large language model (Claude Opus 4.8; Anthropic) was then used to extract structured initial scores for each platform across the ten analytical dimensions, with its reasoning recorded for every score. These initial scores were reviewed and revised by the authors against the primary literature, and a fairness simulation over the full grid of 1,259,712 weight combinations was used to identify and correct structural biases in the recommendation tool. The tool and scoring framework were then deployed as a public beta, and researchers were invited to use the tool and submit feedback through an embedded survey. The resulting researcher responses were re-scored through the revised app (v4.0) and used to validate the recommendations and to refine the scores and algorithm where real-world usage diverged from the simulation (**Figure 4**). The final three stages of this cycle (expert calibration, public deployment, and real-world validation) repeat as new comparative studies are published and additional feedback accrues, with each revision recorded in the calibration history (**Supplementary Note S1**).

#### Platform selection

We included six commercially available plasma proteomics platforms: Olink Explore HT (Olink Proteomics, Uppsala, Sweden), SomaSeq / SomaScan (Illumina, San Diego, CA), NULISA (Alamar Biosciences, Fremont, CA), nELISA/Omni 1000 (Nomic Bio, Montreal, Canada), Proteograph XT (Seer Bio, Redwood City, CA), and TrueDiscovery (Biognosys, Schlieren, Switzerland). We selected these platforms to represent the current landscape of high-throughput protein measurement technologies across affinity-based and mass spectrometry-based approaches. We applied three inclusion criteria: the platform had to be commercially available and actively marketed for biomedical research at the time of writing, support measurement of at least 100 proteins in at least one blood-based matrix, and have at least one peer-reviewed publication demonstrating use in a human cohort study. We excluded platforms in pre-commercial development, legacy platforms with limited ongoing commercial support, and single analyte immunoassay panels. Where a platform technology had evolved substantially across product generations, we scored the most current commercially available configuration. SomaSeq (Illumina) is the NGS successor to the SomaScan microarray assay, briefly marketed as Illumina Protein Prep; most published pQTL and head-to-head evidence cited here derives from the array-based SomaScan readout, as NGS readout validation is still emerging; however, preliminary evidence indicates high correlation between the two products (Spearman *r*=0.991, via). Thus, SomaSeq and SomaScan are treated as essentially equivalent throughout our evaluations. Protein target lists were collected either by downloading directly from the vendor’s websites (Olink, SomaSeq, Alamar), direct correspondence with the vendors (Nomic, Seer), or, in the case of Biognosys TrueDiscovery, from the supplementary materials of Kirsher et al., 2025 (Communications Chemistry, Supplementary Data 8, MS-HAP Depletion arm, 3,575 proteins by primary UniProt accession); we note that the latter is a depletion workflow estimate of protein group counts and should be read as a conservative lower bound relative to particle-enrichment workflows reported elsewhere for the same platform.

#### Scoring framework

We scored each platform across ten performance dimensions on an integer scale from 1 (poor relative to the current platform field) to 5 (best-in-class): proteome coverage, measurement precision, target specificity, sensitivity for low-abundance proteins, cost per sample, throughput, quantification type, sample matrix flexibility, evidence depth, and pQTL accuracy. We report full dimension definitions and scoring criteria in **Supplementary Table S2**. Scores reflect relative standing within the evaluated platform set and do not reflect performance against an absolute external standard.

We assigned scores through structured review of peer-reviewed publications, preprints, and vendor technical documentation, weighting head-to-head comparative studies more heavily than single-platform characterisation reports. Cost scores represented ordinal tiers based on published evidence^24^ and historical quotes provided to the authors for each platform, which varied substantially by sample volume, geography, and commercial agreement. We scored quantification type categorically, assigning higher scores to platforms that provided true absolute quantification (concentration values in physical units traceable to external reference standards) than to those providing relative quantification or normalised signal intensities. We report full platform scores with supporting citations in **Supplementary Tables S3-S9**.

#### Recommendation algorithm (“Help Me Choose”)

The ‘Help Me Choose’ recommendation tool maps user-defined study parameters to a weighted score for each platform, producing a ranked list with explanatory annotations. User inputs spanned three categories, including: (i) study context - i.e., primary research goal from six options, sample matrix type from six options, and estimated study size from four categories, (ii) dimension priorities - i.e., seven sliders each rated on a 1 to 5 scale for cost, proteome coverage, precision, specificity, sensitivity, throughput, and pQTL accuracy, and (iii) mandatory yes/no toggles asking the user if absolute quantification is required, if post-translational modification detection is required, if their study has a central nervous system (CNS) focus, and if their study has a longitudinal / repeat-sampling design. The CNS focus and longitudinal toggles were presented as mandatory questions in the survey interface, ensuring all users engaged with these dimensions and did not leave them at a null default.

Before scoring platforms, the tool applies additional relative weighting that reflects the stated study context. Each research goal adds one to two points to the two or three dimensions it depends on. Cohorts of 1,000 or more add a point to cost and throughput, and cohorts of 10,000 or more add two. A CSF matrix adds four points to sensitivity, and a declared CNS focus adds six to eight more, reflecting generally lower protein concentrations in brain and CSF. These increments are larger than the slider range; accordingly, a CSF study with a CNS focus would surface the ultra-sensitive NULISA assay regardless of what other choices have been made on other dimensions. A platform scoring 1 of 5 on any dimension the user weighted 5 or above would then take a 0.85 multiplier, ensuring strength elsewhere does not mask a critical weakness. Two requirements exclude platforms outright. A strict absolute quantification requirement admits only the Nomic nELISA platform with validated pg/mL concentrations, and a systematic PTM discovery requirement excludes all six assays and recommends orthogonal, specialized methods. A tissue matrix selection does not exclude platforms but drops the sample flexibility score of the three platforms unvalidated for solid tissue to the floor; this penalty is steep enough to be decisive in practice without removing those platforms directly from the ranked list.

Displayed scores are normalized to run from 0 to 100 under the user’s own weights. A platform at the floor of every weighted dimension would map to 0, and one at the ceiling would map to 100. Because platforms cluster in the upper middle of the rubric in real-world settings, evenly weighted priorities produce genuinely similar totals, and displayed gaps widen only as priorities become specific.Most platforms score 3 or 4 on most dimensions, so when a user weights all priorities evenly, the totals come out close together; the gaps between platforms only become large when the user commits to specific priorities. The final production tool therefore groups[1.1] every eligible platform within three calibrated points of the leader into a co-equal top tier and shows a single winner only when one platform separates beyond that three-point band.The production tool therefore reports a co-equal top tier, comprising every eligible platform within three points of the leader, and names a single winner only when the leader is more than three points clear.

##### Effective weight calculation

We derived the effective weight for each scored dimension by combining the user’s slider value with contextual adjustments. Goal-specific increments added +1 to the relevant dimensions, except the biomarker-discovery goal, which added +2 to proteome coverage (**Supplementary Table S11**). For example, drug target identification boosted coverage and specificity; biomarker discovery boosted coverage, throughput, and specificity; disease characterisation and sub-typing boosted coverage, sensitivity, and precision; biomarker validation and clinical translation boosted precision and specificity; pharmacoproteomics (clinical trial-based proteomics) boosted precision and sensitivity; and population-scale proteomics boosted cost and throughput (Supplementary Table S11). Study size applied additive boosts of +0.5 (large cohort: 1,000 to 10,000 samples) or +1 (population-scale: more than 10,000 samples) to the cost and throughput effective weights, capturing the greater practical significance of these factors at scale. Selection of CSF as the sample matrix applied a +4 boost to the sensitivity effective weight, reflecting the low abundance detection requirements of that CNS compartment.

Boost magnitudes were calibrated to ensure that contextually important dimensions exceeded the range of user-adjustable slider values (1 to 5), giving them determinative influence over the platform ranking in scenarios where their relevance was deemed highest-priority. The +4 sensitivity boost for CSF sample type reflects that CSF protein concentrations are approximately 100 to 1,000-fold lower than plasma, making sensitivity the most consequential factor for platform selection in this matrix regardless of the user’s slider setting. The CNS focus boost (sensitivity +6 in non-CSF matrices, +8 in CSF; specificity +2 / +4) reflects the unique analytical demands of CNS proteomics, where detection of low-abundance neurological biomarkers at sub-pg/mL concentrations is a fundamental requirement. Where multiple contextual factors simultaneously elevated sensitivity requirements (e.g., CSF matrix and CNS focus), the corresponding boosts were applied additively, reflecting the independently compounding analytical demands of each factor. The most extreme combination (CSF matrix with CNS focus) correctly identified NULISA as the recommended platform, consistent with its demonstrated attomolar detection of pTau-181, pTau-217, and pTau-231 in CSF. A separate longitudinal / repeat-sampling toggle applied a +2 boost to the precision effective weight, reflecting that repeated-measures and longitudinal study designs place a premium on low between-run variability. All boost magnitudes represent the authors’ calibrated judgements and were not derived from an external published source.

##### Sample matrix flexibility

We applied matrix-specific weights to the Multi-Matrix Validation dimension. Atypical matrixes (tissue, cell culture / screening) or multiple matrices received weight 5, CSF received weight 3, and plasma or serum received weight 1. Within these categories, we applied platform-specific score overrides based on published validation data.

For tissue matrices, we overrode the Multi-Matrix Validation score for Nomic nELISA, NULISA, and Seer Proteograph XT to 1/5, since Nomic nELISA and Alamar NULISA have limited published solid tissue validation, while Seer Proteograph XT’s nanoparticle corona enrichment requires liquid-phase input and is validated for biofluids only. For solid tissue proteomics (including brain tissue), the appropriate Seer product is Proteograph DIRECT, not Proteograph XT.

For cell culture and screening matrices, we overrode the Multi-Matrix Validation score for Nomic nELISA to 4/5 (versus base score 2/5), reflecting extensive published validation in high-throughput cell-based assay contexts^7^.

##### Two-pass scoring

We computed scores in two passes. In the first pass, we calculated a raw score for each platform *p* as:

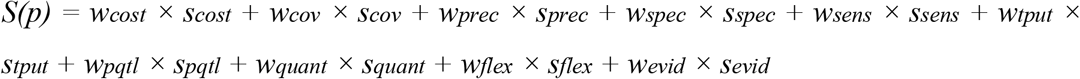

where each *w* represented the effective weight for that dimension and each *s* represented the platform’s score (1 to 5). We set the weight for quantification type (*w_quant*) to 4 when the user required absolute quantification, 1 when preferred but not required, and 0 otherwise. Evidence depth carried a fixed weight of 1 across all configurations. We applied a priority-miss penalty of 0.85 to any platform’s raw score when it scored 1/5 on a dimension the user’s own slider set to 5, or when its Multi-Matrix Validation score for the selected sample type was 1/5, indicating no published validation for that matrix at all. This prevented strong performance on compensatory dimensions from masking a fundamental weakness on the user’s stated priority or, for the matrix case, on a matrix the platform has never been validated for.

In the second pass, we expressed each platform’s raw score as a percentage of the full achievable range for the user’s weighting. For example, a platform scoring 1/5 on every weighted dimension would map to 0% and one scoring 5/5 across the board would map to 100% (displayPct = (S(p) − W·1) / (W·4) × 100, where *W* is the total effective weight). This calibrated scale reports the true separation between platforms; because real platforms cluster in the upper-middle and evenly-weighted priorities yield genuinely similar scores, displayed gaps are often small and widen as priorities become more specific. We additionally grouped every eligible platform within 3 percentage points of the leader into a co-equal “top tier”, presenting a single ranked winner only when one platform separated beyond that band.

##### Hard filters

We marked platforms as ineligible (but not hidden) when they failed a mandatory user requirement, retaining them in the output with explanatory notes. Requiring absolute quantification filtered on capability, excluding every platform whose standard product does not deliver validated absolute quantification in physical concentration units, leaving only Nomic nELISA (ELISA-calibrated pg/mL) eligible among the scored products. Olink Explore HT, SomaSeq, Seer Proteograph XT, Alamar NULISA and Biognosys TrueDiscovery were all excluded; several of these vendors offer absolute quantification through separate products not scored here (Olink Flex / Target 48, Alamar NULISAseq AQ, and Biognosys TrueSignature). Selecting incidental PTM detection retained only DIA-MS platforms. Selecting systematic PTM detection flagged all platforms as ineligible and displayed alternative recommendations, as no evaluated platform offered dedicated PTM enrichment workflows as a standard service.

#### Platform combination tool (“Help Me Combine”)

The platform combination tool, Help Me Combine, ranks pairs of plasma proteomics platforms by how much complementary content they add under user-selected priorities. It operates over all six platforms (Nomic Omni, Olink Explore HT, SomaSeq, Alamar NULISA, Seer Proteograph XT, and Biognosys TrueDiscovery, the two mass-spectrometry platforms scored independently) and therefore over the fifteen distinct pairs.

Each pair was scored on seven axes to help build this combination tool, including: (i) coverage, (ii) complementarity, (iii) combined measurement score, (iv) cost, (v) class balance, (vi) sample volume, and (vii) PTM and proteoform capability. ‘Coverage’ represents the platform pair’s combined coverage on a chosen denominator. ‘Complementarity’ is the net-new content the second platform adds. ‘Combined measurement score’ is defined below; ‘cost’ is the pair’s total cost, while ‘class balance’ measures how evenly the pair spans measurement classes. ‘Sample volume’ is the joint input volume. ‘PTM and proteoform capability’ was set strictly at 1 for any pair containing mass spectrometry and 0 otherwise. Each axis was min-max normalised across the fifteen pairs to the interval 0 to 1, with cost and sample volume inverted so that lower is better. If an axis had zero range across the pairs, it contributed nothing. The composite score for a pair represented the weight-normalised mean of its normalised axes, and the winning pair in any scenario represented the one with the highest composite.

Help Me Combine can be used in two ways. Users who have already run or committed to one of two platforms can name that platform, and the tool ranks the five pairs containing that platform, reporting each pair’s gain over the anchor platform alone. Alternatively, users with no preference or prior selection for either platform can rank all fifteen combinations directly. Users set their weights through a priority picker. By default, baseline axis weights were set at 1 for coverage, complementarity, combined measurement score, and cost, 0.5 for class balance, and 0 for sample volume and PTM/proteoform capability, intended to reflect likely real-world priorities. The user could select up to three of eleven priorities. Five were axis priorities that set their axis weight to 5, including coverage, complementarity, cost, sample volume, and tissue-based work (the latter of which would up-weight mass spectrometry-containing combinations). Six were measurement priorities, including precision, target specificity, sensitivity, absolute quantification, pQTL accuracy, and throughput; selecting one or more would set the combined measurement axis weight to 5 and restrict the score to the selected properties. Weights were never summed across priorities or renormalised outside the composite mean.

The Help Me Combine tool reports uncertainty alongside each ranking. Pair coverage includes bootstrap confidence intervals; leading platforms tied on coverage whose intervals overlap are presented as co-leading; and a per-pair breakdown shows each axis’s share of the composite under the user’s weights, allowing each recommendation to be inspected by the user (**Figure 6B**). Class balance and sample volume are held at reduced or zero default weight to avoid double-counting content the coverage and complementarity axes already capture. Capabilities to detect post-translational modifications and proteoforms are provided as a qualitative flag, since only the two MS platforms (Seer and Biognosys) offer those technical capabilities.

##### Combined measurement score

The combined measurement score is the average of two platforms across whichever measurement properties the user has prioritised. The combined measurement axis is composed of eight individually weightable sub-dimensions, including five intrinsic sub-dimensions (precision, specificity, sensitivity, quantification type, and cis-pQTL) and three maturity sub-dimensions (throughput, evidence depth, and matrix validation). The maturity sub-dimensions default to zero because they reflect differences in maturity and adoption rates between the platforms; newer platforms were not penalised by default and only down-weighted if a user explicitly prioritised platform maturity. As an example, if a user has already committed to Nomic Omni as their first platform and prioritises precision in their choice of paired platform, the tool would score a combination of Nomic (precision = 4) + SomaSeq (precision = 5) as 4.5, Nomic + NULISA (precision = 5) as 4.5, Nomic + Seer (precision = 3) as 3.5, Nomic + Biognosys (precision = 3) as 3.5, or Nomic + Olink (precision = 2) as 3.0. If a user does not select a combined measurement priority, that average runs across all five intrinsic properties (precision, specificity, sensitivity, quantification type, and cis-pQTL accuracy), while the three maturity properties (throughput, evidence depth, and sample flexibility) sit at zero weight until a user selects them. Choosing a measurement priority, such as target specificity, narrows the average to that property alone.

##### Class balance and PTM/proteoform capability

We sought to understand how protein coverage by platform combinations varied across functional protein classes, using the same eleven UniProt-derived classes as the target-class analysis (receptors, kinases, proteases, transporters, ion channels, and related families; **Supplementary Table S10**). To achieve this, we defined “class balance” as how evenly a platform pair’s coverage is distributed across these classes, computed as the Shannon entropy of the class distribution of its combined targets, normalised to the interval 0 to 1. A platform pair detecting many classes in similar proportion scored near 1, while a pair concentrated in a few classes scored lower. Across the fifteen pairs, the axis ranged from 0.77 to 0.93, from the two smallest panels combined (Nomic + Alamar) to the two largest (SomaSeq + Seer Proteograph XT). This axis carried a reduced default weight because the coverage and complementarity axes already captured much of the same content. PTM and proteoform capability were likewise zero-weighted by default and represented the only binary axis, set to 1 for the nine pairs containing a mass-spectrometry platform and 0 for the remaining six, since mass spectrometry is the sole technology in this set that can resolve modifications and proteoforms. It therefore separates pairs into two groups instead of ranking them, and it applies only when a user selects tissue work or PTM detection.

##### Combined coverage and complementarity

We computed coverage and complementarity using the protein lists in the app’s “Protein Coverage” tab, with each platform’s content mapped to UniProt identifiers as described above. Four numeric denominators were used. The druggable target denominator comprised 1,450 unique human proteins that are the mechanism-of-action target of at least one approved or clinical-stage drug, constructed from ChEMBL mechanism records (EBI ChEMBL API^49^): each human single-protein or protein-complex target was mapped to its component gene symbols and annotated with the highest clinical phase of any drug acting on it, giving 929 targets of approved (phase 4) drugs. The FDA-approved biomarker denominator comprised 217 proteins with FDA-approved or - cleared assays, taken from the curated biomarker list distributed with MRMAssayDB^50^ and mapped to UniProt accessions. The detectable proteome union comprised the 15,123 unique proteins measured by at least one of the six platforms, taken directly from the coverage catalogue described above. The canonical proteome denominator comprised the 20,190 reviewed human entries in UniProtKB/Swiss-Prot (organism 9606, reviewed only, accessed July 2026).

For the druggable target and canonical proteome denominators, we computed a pair’s coverage directly by merging the two platforms’ protein lists and counting the unique proteins. For the detectable proteome union, we estimated a pair’s coverage from the two platforms’ individual coverage percentages, assuming their content overlaps at random (coverage of a + coverage of b, minus the expected overlap). Net-new content represented what the pair reaches beyond its stronger member alone. Pair coverage carried bootstrap confidence intervals, and leaders essentially tied on coverage whose intervals overlapped were presented as co-leading instead of being ranked. In the class-level analysis, we assigned each druggable target to a protein class from UniProt annotation, with G-protein-coupled receptors and nuclear receptors identified from the UniProt protein-families field, and computed per-class detectability as a set membership against each platform’s coverage list. Per-class results and multipass membrane counts are in **Supplementary Table S10** and summarised in **Supplementary Figure S4**. We identified targets of specific drug classes from ChEMBL mechanism-of-action records^49^. Molecules were selected by ATC classification (L01E, protein kinase inhibitors, n = 89; L01F, monoclonal antibodies and antibody-drug conjugates, n = 67), with their mechanism records mapped to human target components, and target genes intersected with each platform’s content at the gene level. Mechanism-level target genes included fusion partners (for example BCR via BCR::ABL1); coverage of intracellular targets refers to detection of the circulating protein and does not refer to measurement at the site of drug action.

#### Pathway coverage

We retrieved Reactome pathway membership for all human pathways and mapped to gene symbols, producing 1,285 pathways testable against the canonical proteome. Pathway members that no platform measures were still retained to allow us to compute coverage against the full annotated pathway (restricting the background to the detectable proteome would make every member measurable by construction and inflate coverage). Per-pathway coverage for a platform represented the proportion of that pathway’s members present in the platform’s catalogue, keyed by gene symbol. Coverage efficiency represented that proportion divided by the platform’s panel size in thousands of assays, which normalises for the order-of-magnitude spread in panel size between the focused panels and the discovery-scale catalogues. For the panel of pathways each platform covers most efficiently, pathways were restricted to those with a resolved Reactome name and at least twenty measurable members, and to an efficiency advantage of at least 1.25 times the next-best platform on the same pathway. Pair coverage represents the proportion of pathway members present in the union of the two catalogues, and the pair cost-coverage frontier reports pairs not dominated on both total assays and median coverage across all qualifying pathways.

### Quantification and statistical analysis

#### Fairness validation for the Help Me Choose tool

We assessed whether the ‘Help Me Choose’ recommendation tool systematically advantaged any platform by conducting a simulation across the full user input space. We evaluated all combinations of 6 primary research goals, 6 sample matrix types, 4 study size categories, 3 weight levels (1, 3, and 5) for each of the 7 slider dimensions, and 4 toggle states (CNS focus × longitudinal/repeat-sampling: both off, CNS focus only, longitudinal only, both on), yielding 6 × 6 × 4 × 3⁷ × 4 = 1,259,712 unique scenario-weight combinations. For each combination, we executed the production v4.0 algorithm with the absolute-quantification and PTM toggles set to null and no protein target filtering applied. Rather than record only the top-ranked platform, we identified each scenario’s tie group (i.e., every platform within 3 calibrated points of the leader) to decide whether to display a single winner or a tied set, and scored every member of that group as co-recommended for that scenario. We additionally scored a platform as outright winner in the mutually exclusive subset of scenarios where its tie group had exactly one member.

The fairness simulation’s grid included every study goal, sample type, study size, and priority weight combination, and also incorporated four study characteristic questions set as binary ‘yes/no’ requirements in the user questionnaire, including whether the user required (i) absolute quantification, (ii) PTM detection, (iii) neuroscience-specific assays, or (iv) analysis of longitudinal/repeat samples. Since these inputs are not equally likely in practice, we weighted the grid by their observed prevalence among 63 real-world ‘Help Me Choose’ respondents who completed a beta version of the questionnaire (**Supplementary Table S12**). Under equal weighting, half of all scenarios would be neuroscience-focused, against 7.7% of real respondents; roughly a 6.5-fold overstatement. All results in this paper therefore use the prevalence-weighted grid. An earlier scheme that weighted only the neuroscience and longitudinal questions, leaving study goal, sample type, and study size uniform, is retained for provenance (**Supplementary Table S13**). We evaluated fairness against three criteria: (i) every platform ranks first outright under at least one valid configuration; (ii) every platform is co-recommended in a meaningful share of scenarios; and (iii) each priority slider changes the recommendation when maximised on its own. Scores and weights were refined iteratively until all three of these fairness criteria were met, with the full calibration history provided in **Supplementary Note S1**.

When a respondent marked ‘absolute quantification’ as a required entity, the recommendation tool did not weight scores at all. Instead, it removed every platform whose scored product cannot report physical concentrations, leaving only Nomic nELISA Omni 1000 as the remaining platform to be recommended. Several vendors offer absolute quantification through separate products that we did not score, such as Olink Flex and Target 48, Alamar NULISAseq AQ, and Biognosys TrueSignature. To maintain fairness in scenarios where absolute quantification was required, we highlighted Nomic as the sole mid-to-high-plex recommendation, but listed the lower-plex Olink, Alamar, and Biognosys products as alternative, albeit unassessed, options. When absolute quantitation was preferred but not required, no platform was excluded, and each platform’s graded quantification score (Nomic = 4; Seer, Biognosys = 3; Olink, SomaSeq, NULISA = 2) influenced the rankings instead.

##### Real-world usage validation

To complement the fairness simulation, we analysed recorded usage of the deployed tool from real-world users. From April 2nd 2026 through June 11th 2026, we invited scientists from both co-authors’ networks to complete an early (‘beta’) version of the Help Me Choose questionnaire. We collected 65 study profiles during the validation period, including respondents from academia (46.2%), industry (18.5%), clinical (9.2%) and other (7.7%) backgrounds, ranging from PhD students (18.5%) to post-doctoral researchers (21.5%) to early- (21.5%), mid- (9.2%), and senior-career (23.1%) researchers across the United States (47.7%), United Kingdom (26.2%), France (12.3%), Ireland (6.2%), Spain (3.1%), Netherlands (1.5%), Australia (1.5%) and Germany (1.5%). Their areas of expertise ranged from genomics (24.6%) to clinical biomarkers (15.4%), neuroscience (15.4%), mass spectrometry proteomics (10.8%), computational biology (10.8%), oncology (6.2%), immunology (4.6%), pharmacology (1.5%) and other disciplines (10.8%), with 36.9% of respondents entirely new to proteomics, 38.5% having some experience (1-3 studies), 18.5% regularly running proteomics studies, and 6.2% declaring themselves as experts in proteomics.

During data collection, the recommendation algorithm underwent iterative refinement, meaning users had been served by earlier algorithm versions than the one described in this manuscript (v4.0). To enable a like-for-like comparison, we re-scored all 65 valid profiles through the final algorithm (v4.0, frozen) using the verbatim production computeResults function, excluding a further 2 profiles whose stated requirements (e.g., systematic PTM detection) hard-filtered every platform. This left us with 63 scored profiles. The pooled results reported in the main text of this manuscript are therefore a re-scoring - not the recommendations users observed at the time.

#### Fairness validation for the Help Me Combine tool

We tested whether the Help Me Combine tool structurally favoured any pair of platforms or specific platform by sweeping the production scoring logic across every combination of one to three of the eleven user-selectable priorities (broad coverage, complementary (net-new) targets, precision, target specificity, sensitivity to low-abundance proteins, absolute quantification, pQTL/genetics fit, throughput, low cost, low sample volume, and tissue compatibility - 231 combinations overall), with each evaluated against four coverage denominators (druggable targets (n = 1,450), the detectable-proteome union (n = 15,123), the canonical proteome (n = 20,190), and FDA-approved biomarkers (n = 217)). This produced 924 deterministic scenarios in total. We treated each scenario’s winner as the top composite pair, with no platform fixed in advance and all fifteen pairs of the six platforms eligible. We defined win share as the fraction of scenarios in which a pair was the single top pick, and top 3 share as the fraction in which the pair ranked among the top three. For each platform, we measured the share of scenarios whose winning pair included it. Every winning pair included two platforms, so these shares sum to 200% across the six platforms (not 100%). Because many pairs were statistical ties with overlapping bootstrap confidence intervals, we reported top 3 share alongside win share. Unlike the Help Me Choose tool, no real-world usage data were collected for Help Me Combine; thus, the sweep weighted all priority combinations equally, as no real-world prevalence estimates existed for the different priorities and denominators. Once real selections accumulate, the distribution can be weighted based on prevalence, as was done for the Help Me Choose tool. This fairness simulation is reproducible from the shipped script and tool specification, and the bootstrap intervals use a fixed seed (see **Data and code availability**).

#### Literature growth analysis (outlined in the Discussion section)

We retrieved publication counts from PubMed via the NCBI E-utilities esearch endpoint on 27 July 2026, restricted by publication year. We used three queries: proteomics overall, proteomics[tiab] OR proteomic[tiab]; plasma or serum proteomics in population settings, (plasma[tiab] OR serum[tiab]) AND proteomic*[tiab] AND (cohort[tiab] OR biobank[tiab] OR “population-based“[tiab] OR population-scale[tiab]); and protein quantitative trait studies, pQTL[tiab] OR “protein quantitative trait“[tiab]. We calculated compound annual growth between 2015 and 2025. Absolute counts depended on the query terms chosen; restricting to plasma alone, adding epidemiological terms, or searching all fields rather than title and abstract gave 2015-to-2025 ratios between 7.3 and 13.9 for the population-proteomics series. The tenfold figure quoted in the Discussion should be read as an order-of-magnitude estimate and not as a precise count.

#### Implementation

We built APT as a React 19 application compiled to a self-contained HTML file using Claude Code (Opus 4.8) and deployed it via Vercel. We stored scoring data and platform metadata as version-controlled JSON configuration files, enabling reproducible score updates as new evidence emerges. We deployed the tool at https://aptatlas.org/. The version described here is v4.0 (7 September 2026).

