## Supplementary Materials for "Atlas of Proteomic Technologies: an evidence-based framework for selecting and combining commercial proteomics platforms"

**Supplementary Table S1.** Protein measurement platforms surveyed for the Atlas of Proteomic Technologies (n = 46). The six platforms scored and compared in this manuscript appear first. Integration tier describes how directly a platform's protein list can be compared with others (T1, fixed published per-protein list; T2, defined but narrow or configurable panel; T3, no fixed analyte list, as in open mass spectrometry). Tier describes content list comparability - not platform quality. Availability was checked against vendor documentation in 2026. The full catalogue, with per-platform evidence and citations, is maintained on the “All Platforms” tab of the deployed APT application at. https://aptatlas.org/

| Platform | Vendor | Tier | Status | Note |
|---|---|---|---|---|
| Alamar NULISAseq Inflammation Panel 250 (IP250) | Alamar Biosciences | T1 | Evaluated in this study (combined with Neuro220) | One of the six platforms scored and compared in this manuscript. |
| Alamar NULISAseq Neuro 220 Panel | Alamar Biosciences | T1 | Evaluated in this study (combined with IP250) | One of the six platforms scored and compared in this manuscript. |
| Illumina SomaSeq Discovery (formerly Illumina Protein Prep) | Illumina | T1 | Evaluated in this study (combined with SomaScan) | One of the six platforms scored and compared in this manuscript. |
| Nomic nELISA / Omni 1000 | Nomic Bio | T1 | Evaluated in this study | One of the six platforms scored and compared in this manuscript. |
| Olink Explore HT | Olink (Thermo Fisher Scientific) | T1 | Evaluated in this study | One of the six platforms scored and compared in this manuscript. |
| SomaScan 11K (v5.0) / Illumina SomaScan Discovery | Illumina | T1 | Evaluated in this study (combined with SomaSeq) | One of the six platforms scored and compared in this manuscript. |
| Biognosys TrueDiscovery | Biognosys | T3 | Evaluated in this study | One of the six platforms scored and compared in this manuscript. |
| Seer Proteograph (XT / ONE nanoparticle corona enrichment) | Seer | T3 | Evaluated in this study | One of the six platforms scored and compared in this manuscript. |
| Alamar NULISAseq Immune 340 Panel | Alamar Biosciences | T1 | Candidate for future evaluation | Fixed content list published; not yet scored against the ten dimensions. |
| Alamar NULISAseq Inflammation Panel AQ (absolute quantification) | Alamar Biosciences | T1 | Candidate for future evaluation | Fixed content list published; not yet scored against the ten dimensions. |

| | | | | |
|---|---|---|---|---|
| Olink Reveal | Olink (Thermo Fisher Scientific) | T1 | Candidate for future evaluation | Fixed content list published; not yet scored against the ten dimensions. |
| CellCarta / Caprion targeted MRM & PRM MS biomarker services (incl. Precision Assays immuno-MRM panels) | CellCarta (formerly Caprion Biosciences); immuno-MRM panels via Precision Assays (Paulovich lab, Fred Hutch) acquisition | T2 | Candidate for future evaluation | Fixed content list published; not yet scored against the ten dimensions. |
| MRM Proteomics PeptiQuant Plus / BAK270 targeted MRM kits & services | MRM Proteomics Inc. (UVic-Genome BC Proteomics Centre / Borchers group) | T2 | Candidate for future evaluation | Fixed content list published; not yet scored against the ten dimensions. |
| ProteomEdge targeted-MS panels (ApoEdge, ComplemEdge, LiverEdge, DiscoveryEdge175) | ProteomEdge | T2 | Candidate for future evaluation | Fixed content list published; not yet scored against the ten dimensions. |
| Alamar NULISAqPCR single-plex (e.g. p-tau217) | Alamar Biosciences | T2 | Targeted panel (not proteome-scale) | Defined panel of selected analytes; below the 100-protein threshold or not marketed for proteome-scale discovery. |
| Biognosys TrueSignature (targeted PRM panels) | Biognosys | T2 | Targeted panel (not proteome-scale) | Defined panel of selected analytes; below the 100-protein threshold or not marketed for proteome-scale discovery. |
| Ella Automated Immunoassay System with Simple Plex assays (RUO; Ella CE-IVD variant) | Bio-Techne (ProteinSimple / R&D Systems) | T2 | Targeted panel (not proteome-scale) | Defined panel of selected analytes; below the 100-protein threshold or not marketed for proteome-scale discovery. |

| | | | | |
|---|---|---|---|---|
| Luminex xMAP / INTELLIFLEX bead immunoassay | Luminex/DiaSorin | T2 | Targeted panel (not proteome-scale) | Defined panel of selected analytes; below the 100-protein threshold or not marketed for proteome-scale discovery. |
| Olink Target 48/96 and Flex (custom panels) | Olink (Thermo Fisher Scientific) | T2 | Targeted panel (not proteome-scale) | Defined panel of selected analytes; below the 100-protein threshold or not marketed for proteome-scale discovery. |
| Quanterix Simoa (HD-X / SR-X singleplex and 2-6 plex) | Quanterix | T2 | Targeted panel (not proteome-scale) | Defined panel of selected analytes; below the 100-protein threshold or not marketed for proteome-scale discovery. |
| Targeted MRM/PRM panels (academic and CRO, incl. Inoviv) | Academic core or CRO (e.g. Inoviv) | T2 | Targeted panel (not proteome-scale) | Defined panel of selected analytes; below the 100-protein threshold or not marketed for proteome-scale discovery. |
| Range Biotechnologies translational quantification platform | Range Bio | T3 | Targeted panel (not proteome-scale) | Defined panel of selected analytes; below the 100-protein threshold or not marketed for proteome-scale discovery. |
| Nautilus Voyager (single-molecule iterative mapping) | Nautilus Biotechnology | T2 | Reference entry | No fixed analyte list; included for reference, not panel comparison. |
| Portal Biotech nanopore protein sequencing | Portal Biotech | T3 | Reference entry | No fixed analyte list; included for reference, not panel comparison. |
| Quantum-Si Platinum / Platinum Pro (next-gen protein sequencing) | Quantum-Si | T3 | Reference entry | No fixed analyte list; included for reference, not panel comparison. |
| ProteinXI Human Discovery 400 Panel | ProteinXI (a division of Meso Scale Diagnostics, LLC / MSD) | T1 | Emerging; insufficient published validation | No peer-reviewed human cohort publication at the time of assessment. |

| | | | | |
|---|---|---|---|---|
| Encodia ProteoCode (reverse-translation / DNA-barcoded peptide sequencing) | Encodia, Inc. (San Diego, CA) | T3 | Emerging; insufficient published validation | No peer-reviewed human cohort publication at the time of assessment. |
| Erisyon Fluorosequencing | Erisyon, Inc. (Houston, TX; UT Austin / Marcotte lab technology) | T3 | Emerging; insufficient published validation | No peer-reviewed human cohort publication at the time of assessment. |
| Glyphic Biotechnologies de novo protein sequencing | Glyphic Bio | T3 | Emerging; insufficient published validation | No peer-reviewed human cohort publication at the time of assessment. |
| Primary Bioscience nanopore protein sequencer | Primary Bioscience (Seattle, WA; CoMotion Labs / UW incubator) | T3 | Emerging; insufficient published validation | No peer-reviewed human cohort publication at the time of assessment. |
| Prisma Bio (optical protein sequencing; 'Universal Recognizer') | Prisma Bio / Prisma Tx (founded 2021) | T3 | Emerging; insufficient published validation | No peer-reviewed human cohort publication at the time of assessment. |
| Pumpkinseed deSIPHR nanophotonic single-molecule readout | Pumpkinseed Bio | T3 | Emerging; insufficient published validation | No peer-reviewed human cohort publication at the time of assessment. |
| UNOMR interface nanopore (iNP) / serial nanopore | UNOMR AG (Zurich; ETH Zurich spinout, founded 2023/2025) | T3 | Emerging; insufficient published validation | No peer-reviewed human cohort publication at the time of assessment. |
| Bruker timsTOF (Pro 2 / HT / Ultra 2) with Evosep One | Bruker | T3 | Enabling instrument | Instrument platform on which assay products run; not itself a content-defined assay. |
| Depletion / neat-plasma bottom-up MS (baseline comparator) | N/A, academic / reference method | T3 | Enabling instrument | Instrument platform on which assay products run; not itself a content-defined assay. |
| High-throughput CSF MS resource workflow (5,000-sample scale) | N/A, academic core | T3 | Enabling instrument | Instrument platform on which assay products run; not itself a content-defined assay. |

| | | | | |
|---|---|---|---|---|
| Orbitrap Astral / Astral Zoom (neat or enriched plasma DIA) | Thermo Fisher | T3 | Enabling instrument | Instrument platform on which assay products run; not itself a content-defined assay. |
| SCIEX ZenoTOF 8600 high-throughput DIA | SCIEX (Danaher) | T3 | Enabling instrument | Instrument platform on which assay products run; not itself a content-defined assay. |
| Thermo Scientific Orbitrap Ascend Tribrid (BioPharma / Multi-Omics / Structural Biology editions) | Thermo Fisher Scientific | T3 | Enabling instrument | Instrument platform on which assay products run; not itself a content-defined assay. |
| Thermo Scientific Stellar MS (targeted PRM/MSn instrument line) | Thermo Fisher Scientific | T3 | Enabling instrument | Instrument platform on which assay products run; not itself a content-defined assay. |
| Top-down / intact-proteoform MS (incl. individual-ion MS) | N/A, academic / reference method | T3 | Enabling instrument | Instrument platform on which assay products run; not itself a content-defined assay. |
| Phosphopeptide (PTM) enrichment + Orbitrap Astral or timsTOF | Various (academic / kit) | T3 | Enabling sample preparation | Sample preparation product used upstream of measurement; not itself an assay. |
| PreOmics ENRICH-iST / ENRICHplus (plasma/serum enrichment sample-prep for LC-MS) | PreOmics GmbH (a Bruker company) | T3 | Enabling sample preparation | Sample preparation product used upstream of measurement; not itself an assay. |
| Proteonano Ultraplex Proteomics Platform | Nanomics Biotechnology (Nanomics Biotech) | T3 | Enabling sample preparation | Sample preparation product used upstream of measurement; not itself an assay. |
| Olink Explore 3072 (legacy) | Olink (Thermo Fisher Scientific) | T1 | Retired / superseded | Superseded by Olink Explore HT |
| SomaScan 7K (v4.1) | Standard BioTools → Illumina | T1 | Retired / superseded | Superseded by Illumina SomaScan Discovery (11K, array) or Illumina SomaSeq Discovery (NGS) |

#### Supplementary Table S2. Scoring dimension definitions and criteria

| **Dimension** | **Definition** | **Scoring criteria (1 to 5)** |
|---|---|---|
| Proteome coverage | Number of distinct proteins measurable in a standard workflow | **1:** <1,000 proteins; **2**: 1,000 to 3,000 proteins; **3**: 3,000 to 5,000 proteins; **4**: 5,000 to 7,000 proteins; **5**: >7,000 proteins |
| Precision | Intra- and inter-plate reproducibility; coefficient of variation (CV) | **1:** CV typically >40%; **2**: CV 30 to 40%; **3**: CV 20 to 30%; **4**: CV 10 to 20%; **5**: CV <10% |
| Specificity | Confidence that the measured signal reflects the intended target protein | **1**: high off-target binding risk documented; **2**: moderate risk; **3**: adequate; **4**: high confidence; **5**: comprehensive orthogonal validation of all markers |
| Sensitivity | Detection limit for low-abundance proteins in plasma or CSF | **1**: >1,000 pg/mL (>1 ng/mL); pg/mL range not accessible<br>**2**: 100-1,000 pg/mL; mid-abundance range accessible<br>**3**: 1-100 pg/mL; low pg/mL accessible; moderate sensitivity<br>**4**: 0.01-1 pg/mL; sub-pg/mL detection demonstrated; outperforms standard ELISA<br>**5**: <0.01 pg/mL; attomolar or sub-fg/mL with single-molecule digital counting |
| Cost per sample | Per-sample cost relative to field; higher score = lower cost | **1**: >$3,000/sample; **2**: $1,000 to $3,000; **3**: $250 to $1,000; **4**: $75 to $250; **5**: <$75 |
| Throughput | Sample processing capacity at scale, based on peer reviewed evidence | **1**: <50 samples/day; **3**: moderate throughput; **5**: >1,000 samples/day with automation |
| Quantification type | Absolute vs. relative quantification capability | 1 = qualitative protein readout, only 2 = within-assay relative units only, not comparable across targets (e.g. Olink NPX, SomaScan RFU, NULISA NPQ); **3** = proteome-wide comparable label-free MS intensity (a common scale enabling cross-protein abundance comparison, e.g. iBAQ / Total Protein Approach, but not a physical concentration); **4** = physical concentration in mass units (e.g. ELISA-calibrated pg/mL); **5** = Absolute quantification with isotope-labelled internal standards |

| | | |
|---|---|---|
| Multi-Matrix Validation | Breadth of sample matrices with published validation (plasma, serum, CSF, tissue, cell lysate) | **1**: plasma/serum only; **2**: plasma/serum plus limited validation in one additional matrix; **3**: plasma, serum, and CSF; **4**: plasma, serum, CSF, and one additional matrix with substantial validation; **5**: plasma, serum, CSF, tissue, and cell lysate |
| Evidence depth | Volume and quality of published comparative and large cohort data | **1**: limited preprint data; **3**: several peer-reviewed studies; **5**: multiple large cohort studies and head-to-head comparisons |
| pQTL accuracy | Demonstrated performance in published pQTL studies | **1**: published pQTL studies indicate that this platform is highly unreliable for pQTL detection; **2**: published pQTL studies indicate that this platform is inconsistent for pQTL detection; **3**: published pQTL studies indicate that this platform is inconsistent for pQTL detection but shows strong *cis*-pQTL validation, OR, insufficient published pQTL data to date; **4**: published pQTL validation in at least one cohort showing low risk for epitope effects; **5**: published pQTL validation across multiple cohorts demonstrating consistently low risk for epitope effects. |

***Footnote 1****: Matrix-specific overrides were applied where published evidence diverged from a platform's global Multi-Matrix Validation rating. Nomic nELISA, NULISA, and Seer Proteograph XT were assigned a score of 1 for tissue matrices; Nomic and NULISA had limited published solid tissue validation, and Seer's nanoparticle corona-based enrichment requires liquid-phase sample input, making Proteograph DIRECT or conventional DIA-MS workflows more appropriate for tissue proteomics. Nomic nELISA was assigned a score of 4 for cell culture and screening matrices, reflecting extensive published validation in high-throughput cell-based assay contexts.* ***Footnote 2****: Quantification type scores reflect each vendor's standard high-throughput discovery product. Several evaluated vendors additionally offer absolute quantification (physical pg/mL concentrations) through separate products that are not scored here: Olink (Flex / Target 48 panels with per-assay calibrators), Alamar (NULISAseq AQ panels), and Biognosys (TrueSignature targeted MS).*

**Supplementary Table S3. Overview of platform scores**

| Platform | Coverage | Precision | Specificity | Sensitivity | Cost | Throughput | Quantitation | Multi-matrix flexibility | Evidence | pQTL |
|---|---|---|---|---|---|---|---|---|---|---|
| Olink Explore HT | 4 | 2 | 4 | 4 | 3 | 5 | 2 | 3 | 5 | 3 |
| Illumina SomaSeq Discovery | 5 | 5 | 2 | 4 | 3 | 4 | 2 | 3 | 5 | 2 |
| Alamar NULISA Neuro220 / Inflammation250 | 1 | 5 | 4 | 5 | 4 | 3 | 2 | 3 | 3 | 3 |
| Nomic nELISA Omni 1000 | 2 | 4 | 4 | 3 | 5 | 5 | 4 | 2 | 1 | 3 |
| Seer Proteograph XT | 5 | 3 | 4 | 3 | 3 | 3 | 3 | 3 | 4 | 5 |
| Biognosys TrueDiscovery | 4 | 3 | 4 | 3 | 3 | 3 | 3 | 5 | 3 | 3 |

#### Supplementary Table S4. Olink Explore HT scores and justifications

| Dimension | Score | Justification | Key citation(s) |
|---|---|---|---|
| Proteome coverage | 4 | Covers ~5,400 proteins, placing it second only to SomaSeq among affinity platforms; good breadth but below Seer Proteograph XT and SomaSeq, justifying 4 instead of 5. | *Olink protein list; APT protein coverage tab* Rooney et al., 2025 [1] |
| Precision | 2 | Kirsher et al. (2025) reported a median CV of 26.8% for Olink Explore HT; Rooney et al. (2025) reported an even higher median CV of 35.7%; the mean across these independent cohorts (31.3%) falls within the poor-to-moderate precision range, consistent with a score of 2. | Kirsher et al., 2025 [2] Rooney et al., 2025 [1] |
| Specificity | 4 | PEA requires dual-antibody binding before amplification, substantially reducing off-target signal; Eldjarn et al. 2023 and Sissala et al. 2025 confirm low epitope-artifact rates, though not zero, warranting 4 rather than 5. | Eldjarn et al., 2023 [3] Wik et al., 2021 [4] Sissala et al., 2025 [5] |
| Sensitivity | 4 | PEA's proximity extension amplification step enables sub-pg/mL detection, with Sun et al. 2023 (UKB-PPP) demonstrating measurement of cytokines and tissue-specific proteins below standard ELISA thresholds using Explore 3072, consistent with the 0.01 to 1 pg/mL band that defines Score 4. However, Kirsher et al. 2025 and Rooney et al. 2025 both report that a higher proportion of Explore HT assays fall below the limit of detection in standard plasma than for the earlier Explore 3072 panel, indicating that platform-level sensitivity is uneven across the ~5,400-protein catalogue; the score reflects demonstrated sub-pg/mL capability where detection is achieved, not universal detection across the full panel. | Kirsher et al., 2025 [2] Rooney et al., 2025 [1] Sun et al., 2023 [6] |
| Cost per sample | 3 | Publicly documented Explore HT pricing ranges from $400 to $796 (https://medicine.iu.edu/service-cores/facilities/proteomics/pricing). However, pricing varies substantially, with significant discounts for very large-scale projects such as UKB-PPP. In the only peer-reviewed head-to- | Beimers et al., 2025 [7] |

| | | | |
|---|---|---|---|
| | | head cost comparison, Seer Proteograph XT and Olink Explore HT occupied the same, highest cost tier (“$$$”), with Olink's flat per-sample cost described as comparable to Seer's[7]. | |
| Throughput | 5 | Olink is engineered for large-scale throughput and is proven at population scale. UKB-PPP processed over 54,000 samples in 96-sample batches with automated liquid handling[6]. | Sun et al., 2023 [6] Beimers et al., 2025 [7] |
| Quantification type | 2 | Output in NPX units, which is relative, log-scaled, and anchored to plate controls; no absolute concentration without orthogonal calibration, cross-cohort comparisons require normalization. | Wik et al., 2021 [4] |
| Multi-Matrix Validation | 3 | Validated is primarily in EDTA plasma and serum; several CSF and tissue applications have been reported but with variable performance and matrix-specific dilution requirements. | Sissala et al., 2025 [5] |
| Evidence depth | 5 | Olink, alongside Somalogic, has the largest independent comparative evidence base, including Sissala et al. 2025, Kirsher et al. 2025, and multiple additional head-to-head studies. | Eldjarn et al., 2023 [3] Sissala et al., 2025 [5] Kirsher et al., 2025 [2] |
| pQTL accuracy | 3 | Eldjarn et al. 2023 reports ~72% *cis*-pQTL concordance, which is better than Somalogic. However, residual epitope artifacts, batch normalization effects, and widely reported detection issues for the Explore HT assay constrain the score to 3. | Eldjarn et al., 2023 [3] |

**Supplementary Table S5. Illumina SomaSeq Discovery scores and justifications**

| Dimension | Score | Justification | Key citation(s) |
|---|---|---|---|
| Proteome coverage | 5 | ~9,500 proteins in the current version, the broadest affinity-based catalogue available; longest historical dataset across 5k/7k/11k panel versions. | *SomaSeq protein list; APT protein coverage tab;* Rooney et al., 2025 [1] |
| Precision | 5 | Kirsher et al. 2025 reported a median CV ~5.3% for SomScan; Rooney et al. 2025 reported a median CV of 6.8% for the latest SomScan 11k platform; this is the lowest of all platforms tested, offering best-in-class reproducibility. However, precision metrics may vary for the most recent NGS-based assay. | Rooney et al., 2025 [1]<br>Candia et al., 2022 [8]<br>Candia et al., 2024 [9] |
| Specificity | 2 | Specificity concerns for SomaSeq (whose scores are anchored to the published SomaScan evidence base) are documented primarily through proteogenomic evidence. Eldjarn et al. (2023, Nature; doi:10.1038/s41586-023-06563-x) compared SomaScan v4 in 36,000 Icelanders against Olink Explore 3072 in over 50,000 UK Biobank participants and found that a cis-pQTL, taken as supporting evidence of correct target engagement, was detected for 43% of SomaScan assays versus 72% of Olink assays, despite a similar absolute number on each platform. The same study found that 63% of SomaScan pQTL associations involved variants associated with more than ten proteins, against 52% for Olink, and that 28% of SomaScan proteins with any pQTL had only such non-specific associations, compared with 8% on Olink. | Eldjarn et al., 2023 [3]<br>Joshi and Mayr, 2018 [10]<br>Hoofnagle and MacCoss, 2025 [11] |
| Sensitivity | 4 | SOMAmer reagents achieve femtomolar binding affinities (~$10^{-15}$ M), corresponding to a theoretical detection floor of approximately 0.05 pg/mL for a 50 kDa protein, consistent with the 0.01 to 1 pg/mL Score 4 band. Kirsher et al. 2025 reports 96.2% of proteins detected across all samples with 97% of assays within linear range, confirming broad and consistent low-abundance detection. Note that sub-pg/mL sensitivity figures | Kirsher et al., 2025 [2] |

| | | | |
|---|---|---|---|
| | | should be interpreted alongside the platform's Specificity score of 2/5: some signal at low concentrations may reflect aptamer binding to unintended targets rather than genuine protein detection; these two dimensions are scored independently. | |
| Cost per sample | 3 | Cost per sample for Biognosys, Seer, Olink and SomaSeq/SomaScan varies substantially from the low-to-high hundreds depending on a study's scope and sample size. All four platforms were scored at 3/5, representing par for the field. | Direct quotes received by the authors (see Methods) |
| Throughput | 4 | Supports high-throughput batch processing; SG100K and other large biobanks have run thousands of samples, with an N=50,000 UK Biobank study ongoing, but workflow complexity may introduce slightly more friction than Olink at the largest scales. | Ferkingstad et al., 2021 [12] |
| Quantification type | 2 | Output in relative fluorescence units (RFUs), normalized to reference samples; no absolute quantification, inter-study comparability requires careful normalization. | Candia et al., 2022 [8] |
| Multi-Matrix Validation | 3 | Validated primarily in plasma/serum; several CSF and tissue applications have been published but specificity concerns may be amplified in non-plasma matrices. | Pennington et al., 2026 [13] |
| Evidence depth | 5 | Ferkingstad et al. 2021 (Nature Genetics, deCODE, ~35,000 samples), Eldjarn et al. 2023, Kirsher et al. 2025, and numerous biobank-scale studies have evaluated SomaScan assays; maximum evidence depth alongside Olink. | Ferkingstad et al., 2021 [12]<br>Eldjarn et al., 2023 [3]<br>Kirsher et al., 2025 [2]<br>Rooney et al., 2025 [1] |
| pQTL accuracy | 2 | Over 28,000 pQTLs have been identified using variations of the SomaScan assay, the largest such collection to date and roughly 13,000 more than the closest competitor (Olink). The same limited specificity that lowers target confidence also propagates to genetic analyses: SomaScan's cis-pQTL replication rate (~43%) is lower than the dual-binding affinity platforms, and a greater share of signals reflect aptamer epitope effects rather than the labelled protein, so its pQTL reliability scores below Olink. | Ferkingstad et al., 2021 [12]<br>Eldjarn et al., 2023 [3] |

| | | |
|---|---|---|
| | | Suhre et al. 2025 report broadly similar epitope artifact rates across Olink (~32%) and SomaScan (~31%), suggesting this gap may narrow as further evidence accrues. |

#### Supplementary Table S6. Alamar NULISA scores and justifications

| Dimension | Score | Justification | Key citation(s) |
|---|---|---|---|
| Proteome coverage | 1 | Targeted panels covering ~220 to 500 proteins depending on configuration. Narrowest coverage of all platforms evaluated. | *Alamar protein list; APT protein coverage tab;* Feng et al., 2023 [14] |
| Precision | 5 | NULISA pairs dual antibody capture-and-release with an NGS readout and digital-style quantification, an architecture designed to suppress background and maximise reproducibility. Alamar reports low coefficients of variation across its targeted inflammation and CNS panels, and the focused, heavily optimised panel keeps precision among the best in the comparison for the analytes it measures. | Feng et al., 2023 [14] |
| Specificity | 4 | Sequential dual-antibody capture-and-release requires two independent binding events before a signal is generated, strongly suppressing the single-binder off-target noise that affects aptamer and single-antibody assays. This places NULISA among the most specific affinity platforms, alongside the other dual-binder and mass spectrometry approaches that share the top specificity tier. | Feng et al., 2023 [14] |
| Sensitivity | 5 | Digital proximity ligation with single-molecule counting achieves sub-fg/mL detection for key neurological biomarkers; Shi et al. and Alamar Biosciences validation data report LODs well below 0.01 pg/mL for pTau-181, pTau-217, pTau-231, and NFL, placing NULISA in the attomolar sensitivity tier and clearly warranting the maximum score of 5. | Feng et al., 2023 [14] |
| Cost per sample | 4 | NULISA is typically lower than Olink, Seer, Somalogic or Biognosys due to its targeted panel format and simpler workflow. Its pricing typically stays in the low hundreds per sample – typically not as low as Nomic nELISA, but it remains substantially cheaper than broad high-plex platforms at moderate volumes. | Direct quotes received by the authors |
| Throughput | 3 | NULISA supports moderate throughput in 96-well format; proximity-ligation chemistry involves more handling steps than | Feng et al., 2023 [14] |

| | | | |
|---|---|---|---|
| | | standard affinity platforms, meaning it may be slower at scale than Olink or nELISA. However, several large-scale collaborations between Alamar and global biobanks have been recently announced, such as with the Rhineland study in 23,000 samples and the Global Neurodegeneration Proteomics Consortium in 50,000+ samples; thus, Alamar's throughput score may be increased once publications emerge from these collaborations. | https://alamarbio.com/alamar-biosciences-and-the-german-center-for-neurodegenerative-diseases-dzne-partner-for-landmark-proteomic-profiling-study-in-the-rhineland-study-cohort/<br><br>https://alamarbio.com/proteomics-consortium-profiling-50k-plus-samples-from-patients-with-neurodegenerative-disease/ |
| Quantification type | 2 | NULISA reports NPQ (NULISA Protein Quantification): $\log_2$ normalized counts after within-assay control and interplate normalization. This is a within-assay relative quantification in the same class as Olink NPX or SomaScan RFU. Alamar's separate NULISAseq AQ panels report absolute pg/mL, but that product is not the one scored here. | Feng et al., 2023 [14] |
| Multi-Matrix Validation | 3 | NULISA is validated in plasma and CSF with emerging data in serum and urine. Its dual-antibody format is somewhat more robust to matrix interference than single-binder platforms. | Feng et al., 2023 [14] |
| Evidence depth | 3 | The peer-reviewed evidence base for NULISA is rapidly expanding, but it remains limited relative to Olink and SomaSeq and fewer head-to-head studies or large-scale population cohort studies have been published using the assay as of mid-2026. | Feng et al., 2023 [14] |

| | | | |
|---|---|---|---|
| pQTL accuracy | 3 | Dual-antibody proximity ligation minimizes epitope artifacts that drive spurious pQTLs. However, no published large-scale pQTL dataset exists for NULISA; this score has therefore defaulted to 3/5. | No pQTL studies published to date; defaults to 3/5 |

#### Supplementary Table S7. Nomic nELISA Omni 1000 scores and justifications

| Dimension | Score | Justification | Key citation(s) |
|---|---|---|---|
| Proteome coverage | 2 | The Nomic Omni 1000 panel covers ~1,058 proteins. This is the second narrowest breadth of all platforms assessed. Its current panel does not approach the depth of Olink, SomaSeq, or MS platforms. | *Nomic Omni 1000 protein list;APT protein coverage tab;* Dagher et al., 2025 [15] |
| Precision | 4 | Dagher et al. 2025 reported inter-assay CVs predominantly below 15%. Nomic's bead-based sandwich immunoassay delivers good reproducibility across batches; near best-in-class, though SomaScan's ~5% median CVs have not been independently replicated. | Dagher et al., 2025 [15] |
| Specificity | 4 | Nomic uses a Sandwich ELISA format (two antibodies) providing good specificity; however, cross-reactivity within the highly multiplexed sandwich immunoassay format has not been as extensively characterised as Olink or Alamar. | Dagher et al., 2025 [15] |
| Sensitivity | 3 | Dagher et al. 2025 reported limits of detection of approximately 0.1 pg/mL for representative analytes, which would place selected targets within the 0.01 to 1 pg/mL band. However, this figure derives from a single published study and reflects representative analytes rather than demonstrated panel-wide low-abundance detection, and the microfluidic sandwich-ELISA format lacks the signal-amplification step of proximity-extension or proximity-ligation platforms. Pending independent replication and broader characterisation, Nomic nELISA is scored conservatively at 3 rather than 4; the score will be revisited as further data emerge. | Dagher et al., 2025 [15] |
| Cost per sample | 5 | Nomic Omni 1000 has a widely publicized ~$50/sample list price at moderate to high scale (typically >1,000 samples), representing the lowest-cost high-plex platform available on the market as of mid-2026. | Vendor website, direct quotes received by the authors |
| Throughput | 5 | Nomic processes 1,000+ samples/day using automated microfluidic cartridges. | Dagher et al., 2025 [15] |

| | | | |
|---|---|---|---|
| Quantification type | 4 | nELISA is calibrated to protein standards, enabling semi-quantitative to near-absolute concentration estimates. This is clearly distinguishable from the relative outputs of other affinity-based assays as well as most commercial, high-plex MS assays. | Dagher et al., 2025 [15] |
| Multi-Matrix Validation | 2 | Omni 1000 is validated primarily for plasma and serum; there is limited published evidence for CSF, urine, or tissue lysates. Its proprietary microfluidic cartridge may constrain matrix flexibility. | Dagher et al., 2025 [15] |
| Evidence depth | 1 | Dagher et al. 2025 is the only published validation study; no independent large-scale head-to-head or population cohort deployments have been reported as of early 2026. | Dagher et al., 2025 [15] |
| pQTL accuracy | 3 | nELISA's dual-antibody sandwich format minimizes single-binder epitope artifacts; however, no pQTL study using nELISA has been published yet; thus, this score has defaulted to 3/5. | No pQTL publications to date, hence defaulting to 3/5 |

#### Supplementary Table S8. Seer Proteograph XT scores and justifications

| Dimension | Score | Justification | Key citation(s) |
|---|---|---|---|
| Proteome coverage | 5 | Pietzner et al. 2025 reported ~8,511 protein groups using Seer Proteograph XT. Seer Proteograph ONE, released April 2026, extends this to 10,698 protein groups, representing the broadest proteome coverage in the comparison, on par with or slightly higher than SomaSeq. | Seer Proteograph XT/ONE protein list; APT protein coverage tab;Pietzner et al., 2025 [16] |
| Precision | 3 | Kirsher et al. 2025 reported median CVs of ~26.4%; this is mid-range precision reflecting the inherent variability of DIA-MS on complex proteomes after nanoparticle enrichment. | Kirsher et al., 2025 [2] |
| Specificity | 4 | MS peptide-level identification provides direct molecular evidence of protein identity, eliminating antibody cross-reactivity; however, shared peptides, protein isoforms, and in-source fragmentation can reduce specificity, preventing a score of 5. | Suhre et al., 2025 [17] |
| Sensitivity | 3 | Nanoparticle corona enrichment substantially extends plasma proteome depth relative to unenriched DIA-MS, enabling detection of proteins across an estimated 1 to 100 pg/mL range in plasma; Pietzner et al. 2025 reported approximately 8,500 protein groups detected, including moderately low-abundance species. However, the mass spectrometric detection floor in complex biological matrices constrains absolute sensitivity: very low-abundance proteins below approximately 1 pg/mL are not consistently accessible without orthogonal enrichment, placing Seer below the sub-pg/mL threshold required for a 4/5 score and below the single-molecule sensitivity of NULISA. | Pietzner et al., 2025 [16] |
| Cost per sample | 3 | Cost per sample for Biognosys, Seer, Olink and SomaSeq/SomaScan varies substantially from the low-to-high hundreds depending on a study's scope and sample size. All four platforms were scored at 3/5, representing par for the field. Prior studies have placed Olink Explore HT and Seer Proteograph in the same "$$$" cost tier. | Beimers et al., 2025 [7] |

| | | | |
|---|---|---|---|
| Throughput | 3 | With nanoparticle incubation and a DIA-MS workflow, one machine can process approximately 20 to 40 samples per day; population-scale deployment is feasible, but would require substantial instrument infrastructure. | Beimers et al., 2025 [7] |
| Quantification type | 3 | Label-free DIA-MS yields relative abundances on a proteome-wide comparable intensity scale, supporting cross-protein comparison and approximate intensity-based absolute estimation (e.g. iBAQ / Total Protein Approach), but does not report validated physical concentrations. | Schwanhäusser et al., 2011 [18]<br>Ahrné et al., 2013 [19] |
| Multi-Matrix Validation | 3 | Proteograph is validated in EDTA plasma (Pietzner et al. 2025). The nanoparticle approach can be applied to serum, CSF, and other biofluids, although matrix-specific optimisation is required. | Pietzner et al., 2025 [16] |
| Evidence depth | 4 | Pietzner et al. 2026, Kirsher et al. 2025, and Suhre et al., 2025, provide a growing independent evidence base; good but not yet maximal relative to Olink and Somalogic's publication bases. | Suhre et al., 2025 [17]<br>Pietzner et al., 2025 [16]<br>Kirsher et al., 2025 [2] |
| pQTL accuracy | 5 | Peptide-level MS identification is inherently immune to antibody epitope-binding artifacts that drive spurious pQTLs, measuring peptide sequences rather than aptamer-protein binding events, which eliminates the primary mechanism of pQTL confounding. This has been demonstrated in peer-reviewed studies by Suhre et al., 2025. | Suhre et al., 2025 [17]<br>Pietzner et al., 2025 [16] |

**Supplementary Table S9. Biognosys TrueDiscovery scores and justifications**

| Dimension | Score | Justification | Key citation(s) |
|---|---|---|---|
| Proteome coverage | 4 | DIA-MS with HRM spectral libraries covers ~3,575 (non-enriched) to 7,000 (P2-enriched) protein groups in plasma. This is generally broad MS-based coverage, but it sits below Seer Proteograph XT's nanoparticle-enhanced depth and SomaSeq's 9,500-protein library. | Kirsher (2025) published protein list; APT protein coverage tab;[20] |
| Precision | 3 | Kirsher et al. (2025) reported a median CV of 29.8% for TrueDiscovery. | [2] |
| Specificity | 4 | MS peptide-level detection provides sequence-specific identification inherently more specific than antibody platforms; HRM libraries include retention time calibration and decoy-based FDR control, though shared peptides and isoform ambiguity prevent a score of 5. | [20] |
| Sensitivity | 3 | Standard DIA-MS without nanoparticle or antibody-based pre-enrichment limits detection to the upper approximately 3,000 to 5,000 most abundant plasma proteins, consistent with a 1 to 100 pg/mL sensitivity range. Unlike Seer Proteograph XT, TrueDiscovery does not employ a dedicated low-abundance enrichment step prior to MS acquisition, placing it at the lower end of the Score 3 tier for plasma proteomics; sensitivity improves substantially in less complex matrices such as cell lysates or CSF, where the dynamic range challenge is less extreme. | [20] |
| Cost per sample | 3 | Cost per sample for Biognosys, Seer, Olink and SomaSeq/SomaScan varies substantially from the low-to-high hundreds depending on a study's scope and sample size. All four platforms were scored at 3/5, representing par for the field. | Quotes received directly by the authors and[7] |
| Throughput | 3 | DIA-MS acquisition requires ~60 to 90 minutes instrument time per sample; limits throughput to tens of samples per day per instrument; moderate tier alongside Seer. | [20] |

| | | | |
|---|---|---|---|
| Quantification type | 3 | Label-free DIA-MS produces relative abundances on a proteome-wide comparable intensity scale, supporting cross-protein comparison and approximate intensity-based absolute estimation (e.g. iBAQ / Total Protein Approach), but not validated physical concentrations. Biognosys offers true absolute quantification (pg/mL) via its separate TrueSignature targeted-MS service, which is not the product scored here. | [18]<br>[19] |
| Multi-Matrix Validation | 5 | Validated across plasma, serum, CSF, urine, tissue lysates, cell culture supernatant, and FFPE extracts; this is the widest validated matrix range among the evaluated platforms, reflecting inherent MS versatility. | [20]<br>[21] |
| Evidence depth | 3 | Kirsher et al. 2025 and Sissala et al. 2025 provide independent comparative data; no population-biobank-scale deployment study published, evidence base smaller than Olink, SomaSeq, or Seer. | [2] |
| pQTL accuracy | 3 | No peer-reviewed studies have conducted pQTL discovery using the Biognosys TrueDiscovery platform; thus, this score has defaulted to 3/5. We anticipate an increased score of 4/5 or 5/5 once published proteogenomics studies emerge considering TrueDiscovery is MS-based and presumably less susceptible to epitope effects. | No published pQTL studies to date - hence defaulting to 3/5 |

**Supplementary Table S10.** Class-level coverage of druggable targets across the six platforms. For each of eleven protein classes (assigned by a rule-based UniProt classifier with corrected G-protein-coupled receptor and nuclear-receptor matching), the number of druggable targets (of 1,450) and approved-drug targets (of 929), the broadest single platform and its coverage, the best platform pair and its coverage, the union of all six platforms, and the fraction reachable by any affinity or any mass-spectrometry platform. Data S2 additionally provides the affinity versus mass-spectrometry redundancy partition of the 1,450 druggable targets (292 affinity-only, 152 mass-spectrometry-only, 713 both, 293 detectable by neither) and multipass membrane counts for solute-carrier transporters, ion channels, and G-protein-coupled receptors. Four druggable targets (CTAG1A, IFNA1, HBA1, SMN1) and two approved-drug targets are unclassified by the classifier and are omitted, so class counts sum to 1,446 and 927 rather than 1,450 and 929. Source: ChEMBL druggable target universe and per-platform coverage lists; coverage computed as set membership without per-sample depth adjustment.

| **Class** | **Druggable (n)** | **Approved (n)** | **Broadest single platform** | **%** | **Best pair** | **%** | **Union of six (%)** | **Any affinity (%)** | **Any MS (%)** |
|---|---|---|---|---|---|---|---|---|---|
| Ion channel | 156 | 147 | SomaSeq | 16.0 | SomaSeq + Olink Explore HT | 24.4 | 30.8 | 25.0 | 11.5 |
| GPCR | 154 | 113 | SomaSeq | 26.6 | Seer Proteograph XT + SomaSeq | 40.3 | 47.4 | 34.4 | 20.1 |
| Transcription factor | 24 | 6 | SomaSeq | 58.3 | Seer Proteograph XT + SomaSeq | 70.8 | 75.0 | 66.7 | 45.8 |
| Transporter | 90 | 72 | Seer Proteograph XT | 75.6 | Seer Proteograph XT + SomaSeq | 81.1 | 82.2 | 43.3 | 75.6 |
| Kinase | 142 | 56 | SomaSeq | 71.8 | Seer Proteograph XT + SomaSeq | 82.4 | 85.2 | 77.5 | 69.7 |
| Enzyme (other) | 252 | 185 | Seer Proteograph XT | 69.4 | Seer Proteograph XT + SomaSeq | 86.1 | 88.9 | 73.8 | 72.2 |
| Secreted/ligand | 158 | 77 | SomaSeq | 82.3 | Seer Proteograph XT + SomaSeq | 93.0 | 94.3 | 89.2 | 68.4 |
| Other | 143 | 80 | Seer Proteograph XT | 76.2 | Seer Proteograph XT + SomaSeq | 93.0 | 95.1 | 79.7 | 79.0 |
| Membrane receptor | 192 | 113 | SomaSeq | 92.7 | SomaSeq + Olink Explore HT | 94.8 | 95.3 | 94.8 | 66.7 |
| Protease | 111 | 59 | Seer Proteograph XT | 88.3 | Seer Proteograph XT + SomaSeq | 94.6 | 95.5 | 91.9 | 89.2 |
| Nuclear receptor | 24 | 19 | SomaSeq | 83.3 | Seer Proteograph XT + SomaSeq | 91.7 | 100.0 | 91.7 | 33.3 |

**Supplementary Table S11.** Goal-specific effective weight adjustments applied by the recommendation tool

| Primary goal | Dimensions boosted (magnitude) |
|---|---|
| Drug target identification | Proteome coverage (+1), specificity (+1) |
| Biomarker discovery | Proteome coverage (+2), throughput (+1), specificity (+1) |
| Disease characterisation and sub-typing | Proteome coverage (+1), sensitivity (+1), precision (+1) |
| Biomarker validation and clinical translation | Precision (+1), specificity (+1) |
| Pharmacoproteomics (clinical trials) | Precision (+1), sensitivity (+1) |
| Population proteomics (large biobanks) | Proteome coverage (+1), cost (+1), throughput (+1) |

**Supplementary Table S12.** Prevalence-weighted baseline: co-recommended and outright-win rates from the production v4.0 app over the full 1,259,712-scenario grid, with goal, sample type, study size, and toggle state each reweighted by observed prevalence among 65 real respondents (see Fairness validation, STAR Methods). Co-recommended is the proportion of scenarios in which the platform falls within 3 calibrated points of the leader (its tie group has two or more members); outright win is the proportion in which its tie group has exactly one member. Rows sorted by canonical proteome coverage.

| Platform | Co-recommended | Outright win |
|---|---|---|
| Seer Proteograph XT | 39.1% | 12.3% |
| SomaSeq | 33.4% | 9.4% |
| Olink Explore HT | 33.2% | 3.1% |
| Biognosys TrueDiscovery | 16.5% | 1.0% |
| Nomic nELISA | 28.3% | 6.7% |
| Alamar NULISA | 17.4% | 4.7% |

**Supplementary Table S13.** Superseded reference, retained for provenance; the current baseline is shown in Supplementary Table S12. Outright win-shares under the old v2.5 methodology across the full grid of 1,259,712 scenario-weight combinations, weighted by the prevalence estimates in use at the time (CNS focus at ~20% of sessions, later corrected to 7.7%; longitudinal at an imputed 17.5%). The v2.5 tally recorded a single winning platform per scenario and no co-recommendation statistic, so the six shares sum to 100%. The current methodology recognises co-recommended ties, so its outright rates are lower for every platform and are not directly comparable: Olink Explore HT falls from 18.2% here to 3.1% in Table S12, Alamar NULISA from 19.5% to 4.7%, and Biognosys TrueDiscovery shows the largest proportional change, from 13.5% to 1.0%. Ranks also shift by up to three positions: Seer Proteograph XT stands fourth here but first in Table S10, and Nomic nELISA sixth here but third.

| **Platform** | **Outright win-share** |
|---|---|
| SomaSeq | 22.7% |
| Alamar NULISA | 19.5% |
| Olink Explore HT | 18.2% |
| Seer Proteograph XT | 14.6% |
| Biognosys TrueDiscovery | 13.5% |
| Nomic nELISA Omni 1000 | 11.6% |

**Supplementary Table S14.** Help Me Combine priority sweep: per-pair win-share and top-3 share across all fifteen pairs of the six coverage-bearing platforms (Seer Proteograph XT and Biognosys TrueDiscovery scored as independent mass-spectrometry platforms), 924 deterministic scenarios (231 priority combinations x 4 coverage denominators).

| **Platform pair** | **Win %** | **Top 3 %** |
|---|---|---|
| Nomic Omni + Seer Proteograph XT | 18.7 | 48.5 |
| Nomic Omni + Olink Explore HT | 15.6 | 33.4 |
| SomaSeq + Seer Proteograph XT | 13.3 | 28.7 |
| Olink Explore HT + Seer Proteograph XT | 12.1 | 31.8 |
| Nomic Omni + Alamar NULISA | 8.9 | 23.5 |
| Alamar NULISA + Seer Proteograph XT | 6.3 | 20.1 |
| SomaSeq + Alamar NULISA | 6.3 | 14.0 |
| Olink Explore HT + Biognosys TrueDiscovery | 5.0 | 18.7 |
| Nomic Omni + Biognosys TrueDiscovery | 4.9 | 26.0 |
| Nomic Omni + SomaSeq | 4.3 | 22.2 |
| Olink Explore HT + SomaSeq | 2.2 | 11.6 |
| Olink Explore HT + Alamar NULISA | 1.7 | 9.4 |
| Alamar NULISA + Biognosys TrueDiscovery | 0.5 | 5.4 |
| SomaSeq + Biognosys TrueDiscovery | 0.2 | 3.1 |
| Seer Proteograph XT + Biognosys TrueDiscovery | 0.0 | 3.6 |

*Outright-win counts and shares displayed to users at the moment of submission, split by which tool version was live when each profile was submitted (earlier tool versions, n = 47; a later development version, n = 18). Counts are given alongside percentages because the later-version cohort is small: one profile is 5.6% of it, so differences of a single profile should not be read as trends. The two columns are not a before-and-after comparison on a fixed sample: they describe different profiles served by different tool versions, so any difference between columns confounds the algorithm change with the change in cohort. For a like-for-like comparison, every collected profile is re-scored through the single final tool in Supplementary Figure S1, and this table exists only to document what users were actually shown during development, which the re-scored figures deliberately do not report.*

**Supplementary Table S15. FDA-approved or -cleared protein biomarkers (n = 217) and their measurability on each of the six evaluated platforms.** Platforms column gives the number of platforms measuring each biomarker. Rows are ordered by decreasing platform count, then gene symbol. Biomarker list derived from the curated set distributed with MRMAssayDB (Bhowmick et al. 2021), deduplicated to unique UniProt accessions. Three biomarkers (ACP4, EGR1, LDHAL6A) are measured by no platform. Per-platform totals: Nomic Omni 90, Alamar NULISA 31, Olink Explore HT 125, SomaSeq 195, Seer Proteograph XT 187, Biognosys TrueDiscovery 158.

| Gene | UniProt | Platforms | Nomic Omni | Alamar NULISA | Olink Explore HT | SomaSeq | Seer Proteograph XT | Biognosys True Discovery |
|---|---|---|---|---|---|---|---|---|
| ACHE | P22303 | 6 | Y | Y | Y | Y | Y | Y |
| APOA1 | P02647 | 6 | Y | Y | Y | Y | Y | Y |
| APOE | P02649 | 6 | Y | Y | Y | Y | Y | Y |
| APOH | P02749 | 6 | Y | Y | Y | Y | Y | Y |
| CST3 | P01034 | 6 | Y | Y | Y | Y | Y | Y |
| GOT1 | P17174 | 6 | Y | Y | Y | Y | Y | Y |
| IL1RL1 | Q01638 | 6 | Y | Y | Y | Y | Y | Y |
| MMP9 | P14780 | 6 | Y | Y | Y | Y | Y | Y |
| MPO | P05164 | 6 | Y | Y | Y | Y | Y | Y |
| PDGFRB | P09619 | 6 | Y | Y | Y | Y | Y | Y |
| PPBP | P02775 | 6 | Y | Y | Y | Y | Y | Y |
| ACP1 | P24666 | 5 | Y |  | Y | Y | Y | Y |
| ACP3 | P15309 | 5 | Y |  | Y | Y | Y | Y |
| ACP5 | P13686 | 5 | Y |  | Y | Y | Y | Y |
| AFP | P02771 | 5 | Y |  | Y | Y | Y | Y |
| AGT | P01019 | 5 | Y |  | Y | Y | Y | Y |
| AHSG | P02765 | 5 | Y |  | Y | Y | Y | Y |
| BCHE | P06276 | 5 | Y |  | Y | Y | Y | Y |
| C1QA | P02745 | 5 |  | Y | Y | Y | Y | Y |
| C3 | P01024 | 5 | Y |  | Y | Y | Y | Y |
| C4B | P0C0L5 | 5 | Y |  | Y | Y | Y | Y |
| C5 | P01031 | 5 | Y |  | Y | Y | Y | Y |
| CD274 | Q9NZQ7 | 5 | Y | Y | Y | Y | Y |  |
| CD36 | P16671 | 5 | Y |  | Y | Y | Y | Y |
| CD4 | P01730 | 5 | Y | Y | Y | Y | Y |  |
| CKB | P12277 | 5 | Y |  | Y | Y | Y | Y |
| CRP | P02741 | 5 | Y | Y |  | Y | Y | Y |
| F10 | P00742 | 5 | Y |  | Y | Y | Y | Y |
| F11 | P03951 | 5 | Y |  | Y | Y | Y | Y |
| F2 | P00734 | 5 | Y |  | Y | Y | Y | Y |
| F7 | P08709 | 5 | Y |  | Y | Y | Y | Y |
| F9 | P00740 | 5 | Y |  | Y | Y | Y | Y |
| FN1 | P02751 | 5 | Y |  | Y | Y | Y | Y |
| FTL | P02792 | 5 | Y |  | Y | Y | Y | Y |
| GBA | P04062 | 5 | Y |  | Y | Y | Y | Y |
| GFAP | P14136 | 5 | Y | Y | Y | Y | Y |  |
| GH1 | P01241 | 5 | Y |  | Y | Y | Y | Y |
| GP5 | P40197 | 5 | Y |  | Y | Y | Y | Y |
| IDUA | P35475 | 5 | Y |  | Y | Y | Y | Y |
| IGFBP1 | P08833 | 5 | Y |  | Y | Y | Y | Y |
| IGFBP3 | P17936 | 5 | Y |  | Y | Y | Y | Y |
| IL2RA | P01589 | 5 | Y | Y | Y | Y |  | Y |
| IL2RG | P31785 | 5 | Y |  | Y | Y | Y | Y |
| LGALS3 | P17931 | 5 | Y | Y | Y |  | Y | Y |
| MUC16 | Q8WXI7 | 5 | Y | Y | Y |  | Y | Y |

| | | | | | | | | |
|---|---|---|---|---|---|---|---|---|
| NPPB | P16860 | 5 | Y | Y | Y | Y | Y | |
| PF4 | P02776 | 5 | Y | | Y | Y | Y | Y |
| PLAU | P00749 | 5 | Y | | Y | Y | Y | Y |
| PROC | P04070 | 5 | Y | | Y | Y | Y | Y |
| PROS1 | P07225 | 5 | Y | Y | | Y | Y | Y |
| PRSS2 | P07478 | 5 | Y | | Y | Y | Y | Y |
| RBP4 | P02753 | 5 | Y | | Y | Y | Y | Y |
| S100A9 | P06702 | 5 | Y | Y | | Y | Y | Y |
| SERPINA1 | P01009 | 5 | Y | | Y | Y | Y | Y |
| SERPINC1 | P01008 | 5 | Y | | Y | Y | Y | Y |
| SERPINE1 | P05121 | 5 | Y | | Y | Y | Y | Y |
| SERPINF2 | P08697 | 5 | Y | | Y | Y | Y | Y |
| SHBG | P04278 | 5 | Y | | Y | Y | Y | Y |
| SORD | Q00796 | 5 | Y | | Y | Y | Y | Y |
| TF | P02787 | 5 | Y | | Y | Y | Y | Y |
| TFRC | P02786 | 5 | Y | | Y | Y | Y | Y |
| VWF | P04275 | 5 | Y | | Y | Y | Y | Y |
| A2M | P01023 | 4 | Y | | | Y | Y | Y |
| ACE | P12821 | 4 | Y | | Y | | Y | Y |
| ACP6 | Q9NPH0 | 4 | | | Y | Y | Y | Y |
| ALB | P02768 | 4 | Y | | | Y | Y | Y |
| ALPI | P09923 | 4 | | | Y | Y | Y | Y |
| AMBP | P02760 | 4 | | | Y | Y | Y | Y |
| APOB | P04114 | 4 | | | Y | Y | Y | Y |
| B2M | P61769 | 4 | | | Y | Y | Y | Y |
| BTD | P43251 | 4 | | | Y | Y | Y | Y |
| C4A | P0C0L4 | 4 | Y | | | Y | Y | Y |
| CAT | P04040 | 4 | | | Y | Y | Y | Y |
| CKM | P06732 | 4 | Y | | | Y | Y | Y |
| CKMT1A | P12532 | 4 | | | Y | Y | Y | Y |
| ERBB2 | P04626 | 4 | Y | | Y | Y | | Y |
| F12 | P00748 | 4 | | | Y | Y | Y | Y |
| F13B | P05160 | 4 | | | Y | Y | Y | Y |
| F8 | P00451 | 4 | Y | | | Y | Y | Y |
| FGA | P02671 | 4 | | | Y | Y | Y | Y |
| FTH1 | P02794 | 4 | | Y | | Y | Y | Y |
| GLA | P06280 | 4 | Y | | | Y | Y | Y |
| GOT2 | P00505 | 4 | Y | | | Y | Y | Y |
| GP1BA | P07359 | 4 | | | Y | Y | Y | Y |
| GP1BB | P13224 | 4 | | | Y | Y | Y | Y |
| GPI | P06744 | 4 | | | Y | Y | Y | Y |
| GPT | P24298 | 4 | Y | | | Y | Y | Y |
| GSR | P00390 | 4 | | | Y | Y | Y | Y |
| GUSB | P08236 | 4 | | | Y | Y | Y | Y |
| HBA1 | P69905 | 4 | | Y | | Y | Y | Y |
| HLA-A | P04439 | 4 | | | Y | Y | Y | Y |
| HLA-DRA | P01903 | 4 | | Y | Y | | Y | Y |
| HP | P00738 | 4 | Y | | | Y | Y | Y |
| IDH1 | O75874 | 4 | Y | | | Y | Y | Y |
| IGF1 | P05019 | 4 | Y | | | Y | Y | Y |
| ITGB1 | P05556 | 4 | | | Y | Y | Y | Y |
| KLKB1 | P03952 | 4 | | | Y | Y | Y | Y |
| KRT19 | P08727 | 4 | | | Y | Y | Y | Y |
| LAP3 | P28838 | 4 | | | Y | Y | Y | Y |
| LHB | P01229 | 4 | Y | | Y | Y | | Y |
| LPA | P08519 | 4 | | | Y | Y | Y | Y |
| LTF | P02788 | 4 | | | Y | Y | Y | Y |
| MB | P02144 | 4 | Y | | | Y | Y | Y |

| | | | | | | | | |
|---|---|---|---|---|---|---|---|---|
| MUC1 | P15941 | 4 | Y | | Y | Y | Y | |
| NT5E | P21589 | 4 | | | Y | Y | Y | Y |
| ORM1 | P02763 | 4 | Y | | Y | | Y | Y |
| PKLR | P30613 | 4 | | | Y | Y | Y | Y |
| PLA2G7 | Q13093 | 4 | | | Y | Y | Y | Y |
| PLG | P00747 | 4 | | | Y | Y | Y | Y |
| PNLIP | P16233 | 4 | | | Y | Y | Y | Y |
| PRL | P01236 | 4 | | | Y | Y | Y | Y |
| PRSS1 | P07477 | 4 | Y | | | Y | Y | Y |
| PRTN3 | P24158 | 4 | | | Y | Y | Y | Y |
| RBP7 | Q96R05 | 4 | | | Y | Y | Y | Y |
| REN | P00797 | 4 | Y | | Y | Y | | Y |
| S100A8 | P05109 | 4 | Y | | | Y | Y | Y |
| SERPING1 | P05155 | 4 | Y | | Y | Y | | Y |
| TCN1 | P20061 | 4 | | | Y | Y | Y | Y |
| TG | P01266 | 4 | | | Y | Y | Y | Y |
| TGM2 | P21980 | 4 | | | Y | Y | Y | Y |
| TNFRSF8 | P28908 | 4 | Y | Y | Y | Y | | |
| TNNT2 | P45379 | 4 | Y | | Y | Y | Y | |
| TPMT | P51580 | 4 | | | Y | Y | Y | Y |
| ACP2 | P11117 | 3 | | | | Y | Y | Y |
| ALDOA | P04075 | 3 | | | | Y | Y | Y |
| ALDOB | P05062 | 3 | | | | Y | Y | Y |
| ALDOC | P09972 | 3 | | | | Y | Y | Y |
| ALK | Q9UM73 | 3 | Y | | Y | Y | | |
| ALPG | P10696 | 3 | | | Y | Y | | Y |
| ALPL | P05186 | 3 | | | | Y | Y | Y |
| BGLAP | P02818 | 3 | | | Y | Y | Y | |
| C1QB | P02746 | 3 | | | | Y | Y | Y |
| C1QC | P02747 | 3 | | | | Y | Y | Y |
| CALCA | P01258 | 3 | Y | | Y | | Y | |
| CGA | P01215 | 3 | | | | Y | Y | Y |
| CP | P00450 | 3 | | | | Y | Y | Y |
| EPO | P01588 | 3 | | Y | Y | Y | | |
| F13A1 | P00488 | 3 | | | | Y | Y | Y |
| F5 | P12259 | 3 | | | | Y | Y | Y |
| FGB | P02675 | 3 | | | | Y | Y | Y |
| G6PD | P11413 | 3 | | | | Y | Y | Y |
| GAA | P10253 | 3 | | | | Y | Y | Y |
| GGT1 | P19440 | 3 | | | Y | | Y | Y |
| GP9 | P14770 | 3 | | | | Y | Y | Y |
| HBB | P68871 | 3 | | | | Y | Y | Y |
| HLA-B | P01889 | 3 | | | | Y | Y | Y |
| HLA-DQB1 | P01920 | 3 | | | | Y | Y | Y |
| HPX | P02790 | 3 | | | | Y | Y | Y |
| IDH2 | P48735 | 3 | | | | Y | Y | Y |
| IGF2 | P01344 | 3 | | | | Y | Y | Y |
| IL2RB | P14784 | 3 | | Y | Y | Y | | |
| ITGA2 | P17301 | 3 | | | | Y | Y | Y |
| ITGA2B | P08514 | 3 | | | | Y | Y | Y |
| ITGB3 | P05106 | 3 | | | | Y | Y | Y |
| KRAS | P01116 | 3 | Y | | | Y | Y | |
| LDHA | P00338 | 3 | | | | Y | Y | Y |
| LDHB | P07195 | 3 | | | | Y | Y | Y |
| LRRK2 | Q5S007 | 3 | | Y | | Y | Y | |
| LYZ | P61626 | 3 | | | | Y | Y | Y |
| ORM2 | P19652 | 3 | | | | Y | Y | Y |
| PKM | P14618 | 3 | | | | Y | Y | Y |

| | | | | | | | | |
|---|---|---|---|---|---|---|---|---|
| PLAT | P00750 | 3 | | | | Y | Y | Y |
| POMC | P01189 | 3 | | | Y | Y | Y | |
| PRSS3 | P35030 | 3 | | | | Y | Y | Y |
| PTH | P01270 | 3 | Y | | Y | Y | | |
| PTPRC | P08575 | 3 | | | Y | | Y | Y |
| PTPRN | Q16849 | 3 | | | Y | Y | Y | |
| RBP1 | P09455 | 3 | | | Y | Y | Y | |
| RBP2 | P50120 | 3 | | | Y | Y | Y | |
| RBP5 | P82980 | 3 | | | Y | Y | Y | |
| SSB | P05455 | 3 | | | Y | Y | Y | |
| TNNI3 | P19429 | 3 | | | Y | Y | Y | |
| TOP2A | P11388 | 3 | Y | | | Y | Y | |
| TTR | P02766 | 3 | | | Y | | Y | Y |
| UCHL1 | P09936 | 3 | | Y | | Y | Y | |
| WFDC2 | Q14508 | 3 | | | Y | Y | Y | |
| ACP7 | Q6ZNF0 | 2 | | | | Y | Y | |
| AMY2A | P04746 | 2 | | | | Y | | Y |
| BRAF | P15056 | 2 | | | | Y | Y | |
| CALCA | P06881 | 2 | | Y | | Y | | |
| CKMT2 | P17540 | 2 | | | | Y | Y | |
| CYP2D6 | P10635 | 2 | | | | Y | Y | |
| CYP3A5 | P20815 | 2 | | | | Y | Y | |
| DPYD | Q12882 | 2 | | | | | Y | Y |
| GALC | P54803 | 2 | | | | Y | Y | |
| GALT | P07902 | 2 | | | | | Y | Y |
| GAST | P01350 | 2 | | | Y | | Y | |
| HLA-DQA1 | P01909 | 2 | | | | Y | Y | |
| HLA-DRB1 | P01911 | 2 | | | | | Y | Y |
| HLA-DRB3 | P79483 | 2 | | | | Y | Y | |
| HLA-DRB4 | P13762 | 2 | | | | Y | Y | |
| HLA-DRB5 | Q30154 | 2 | | | | | Y | Y |
| KLK3 | P07288 | 2 | Y | | | Y | | |
| LDHAL6B | Q9BYZ2 | 2 | | | | Y | Y | |
| MSH2 | P43246 | 2 | | | | Y | Y | |
| MSH6 | P52701 | 2 | | | | Y | Y | |
| NUMA1 | Q14980 | 2 | | | | Y | Y | |
| TFR2 | Q9UP52 | 2 | | | | Y | | Y |
| TSHB | P01222 | 2 | | | Y | Y | | |
| UGT1A1 | P22309 | 2 | | | | Y | Y | |
| AMHR2 | Q16671 | 1 | | | | Y | | |
| AMY2B | P19961 | 1 | | | | Y | | |
| FTMT | Q8N4E7 | 1 | | | | Y | | |
| GOT1L1 | Q8NHS2 | 1 | | | | Y | | |
| INHA | P05111 | 1 | | | | Y | | |
| LDHC | P07864 | 1 | | | | Y | | |
| MLH1 | P40692 | 1 | | | | Y | | |
| MTHFR | P42898 | 1 | | | | | | Y |
| PMS2 | P54278 | 1 | | | | Y | | |
| RBP3 | P10745 | 1 | | | | Y | | |
| RHD | Q02161 | 1 | | | | | Y | |
| SLCO1B1 | Q9Y6L6 | 1 | | | | | Y | |
| TPSD1 | Q9BZJ3 | 1 | | | Y | | | |
| VKORC1 | Q9BQB6 | 1 | | | | | Y | |
| ACP4 | Q9BZG2 | 0 | | | | | | |
| EGR1 | P18146 | 0 | | | | | | |
| LDHAL6A | Q6ZMR3 | 0 | | | | | | |

170

#### Supplementary Note S1. Algorithm calibration history

We refined the recommendation algorithm through thirteen iterative calibration steps following initial implementation, each motivated by an identified fairness or accuracy consideration. Steps 1 to 9 preceded the v3.0 re-validation; steps 10 to 12 record its findings, and step 13 records the subsequent v3.5 evidence review. Every step is dated to a tool version and reproduced in the version history published with the application, so the two records can be checked against each other. A user-facing change log for the deployed application, including UI and documentation releases not relevant to scoring, is maintained at aptatlas.org (Methods tab); this note records the methodological calibration history only, with steps labelled by the public release that shipped them.

**1**. Olink/SomaSeq win-share imbalance corrected. Initial simulation runs showed Olink winning 4.8 percentage points more scenarios than SomaSeq. We identified that the specificity dimension received goal-specific boosts from three goals (drug target identification, biomarker discovery, and biomarker validation) versus precision's two boosts (biomarker validation and pharmacoproteomics). We added a +1 precision boost to the disease characterisation goal, which brought the gap within 1 percentage point and better reflected the measurement fidelity requirements of disease sub-typing studies.

**2.** Score compression revisited. Initial percentage scores clustered in the 65 to 87% range, making differences difficult to interpret. We first replaced normalisation against the theoretical maximum (total weight × 5) with normalisation against the highest-scoring platform in each user configuration (winner = 100%). This was subsequently found to overstate separation and was replaced with an anchored, calibrated scale (step 8); the original compressed band, in retrospect, described the honest scale now used by design.

**3.** Priority-miss penalty introduced. Platforms with a score of 1/5 on a dimension the user had flagged as high priority (effective weight ≥5) were receiving artificially inflated rankings due to strong scores on other dimensions. We applied a 0.85× multiplicative penalty in these cases. In practice, this fired most often for Nomic nELISA and NULISA when proteome coverage was weighted highly, correctly suppressing them in coverage-prioritised discovery contexts.

**4.** Absolute quantification weight reduced. A "nice to have" absolute quantification response was initially weighted at 2, giving Nomic a roughly 5 percentage point advantage in scenarios where users expressed only mild interest in absolute quantification. We reduced the nice-to-have weight to 1, reserving the stronger weight (4) for users who explicitly required absolute quantification.

**5.** Cell lysates replaced with cell culture/screening. The original sample type "cell lysates" did not capture the perturb-seq and high-throughput cell-based screening use case for which Nomic nELISA has been most extensively validated. We replaced this option with "cell culture/screening" and applied a Multi-Matrix Validation override of 4/5 for Nomic in this context (versus base score 2/5), reflecting published validation in perturb-seq and drug screening workflows.

**6.** Seer Proteograph XT tissue penalty applied. Seer Proteograph XT received a base Multi-Matrix Validation score of 3/5, matching Olink and SomaSeq. However, its nanoparticle corona depletion technology requires liquid-phase sample input and has not been validated for solid tissue proteomics. We applied an override of 1/5 for the tissue sample type, matching the penalty applied to Nomic and NULISA. For solid tissue workflows, the appropriate Seer product is Proteograph DIRECT, not Proteograph XT.

**7.** Olink and SomaSeq tissue homogenate penalty applied. Following the Seer correction, Olink Explore HT ranked first in tissue scenarios due to its strong specificity and sensitivity scores. However, both Olink and SomaSeq are validated in tissue homogenates only, not in native solid tissue MS workflows. We applied an override of 2/5 for both platforms in tissue scenarios, below their base score of 3/5 but above the 1/5 assigned to platforms with no tissue validation at all. TrueDiscovery (Biognosys) retained its base score of 5/5, reflecting its native DIA-MS workflow that handles digested solid tissue from any matrix including brain. This override was later removed after an evidence review (step 13).

**8.** Calibrated, anchored scale and top-tier tie grouping introduced (v2.3). Winner-normalisation (step 2) was subsequently found to overstate confidence, displaying a median ~2-point separation as a decisive-looking lead and pinning the leader to 100% in 17 of the 18 responses collected under that scheme. We replaced it with an anchored, calibrated scale, each platform's score expressed on a 0 to 100% range where 1/5 on every weighted dimension maps to 0% and 5/5 across the board maps to 100%, and grouped every eligible platform within 3 calibrated points of the leader into a co-equal top tier, reporting a sole winner only when one platform separated beyond that band. In the same revision we removed the cohort-comparability display bonus, which was an unreachable code path in the deployed tool that never affected a live recommendation. Platform rankings were unchanged by these revisions; only the displayed magnitude of separation and the single-winner-versus-tie presentation changed.

**9.** Quantification handling corrected (v2.4). The 'absolute quantification required' hard filter previously excluded only platforms scoring below 3/5 on quantification type, which admitted semi-quantitative platforms whose standard output is not a physical concentration, notably Seer Proteograph XT (label-free DIA). We replaced the score threshold with a capability test: an absolute-quantification requirement now admits only platforms whose scored standard product reports validated physical concentration output, which among the evaluated products is Nomic nELISA alone (ELISA-calibrated pg/mL). We also clarified the quantification-type axis, 4 = physical concentration; 3 = proteome-wide comparable label-free MS intensity; 2 = within-assay relative units, and moved Alamar NULISA from 3 to 2 to match its NPQ output (log2 normalised counts, within-assay relative, the same class as Olink NPX). This was the only platform score changed, and no weights changed. Where a vendor offers absolute quantification through a separate product not scored here (Olink Flex / Target 48, Alamar NULISAseq AQ, Biognosys TrueSignature), this is noted. Platform rankings were otherwise unchanged.

**10.** Fairness simulation methodology corrected (v3.0). A full re-validation of the fairness simulation against the current, unchanged score matrix identified four issues, none of which involved any platform's score. Two were in how the simulation tallied a win rather than in the recommendation tool itself, and are described here; the remaining two are steps 11 and 12. From v2.5 onward, every reported win-share figure is reproducible from the archived tool specification and sweep script, and the v2.5 figures are retained in Supplementary Table S11; figures displayed during earlier development were not version-pinned and are superseded. First, win had been defined as whichever platform had the single highest score, even when its margin over the next platform was under the app's own 3-point tie threshold; the app itself displays such cases as a statistical tie, not a sole winner, so the simulation's tallying disagreed with what users actually see. Win is now defined identically to the app's own condition: a platform is an outright winner only when its tie group (every platform within 3 calibrated points of the leader) has exactly one member;

all other tie-group members are scored as co-recommended, a separate, non-exclusive statistic reported alongside outright win throughout. Second, the earlier simulation had treated every goal, sample type, study size, and toggle-state combination as equally likely. The baseline is now reweighted by observed prevalence among 65 real respondents across all four axes, replacing an earlier version that reweighted only the two toggles. Under the corrected methodology, Seer Proteograph XT led the simulated baseline on both statistics at that step (38.3% co-recommended, 13.5% outright), rather than SomaSeq under the previous uniform, win-only tally. The previous methodology is retained, unchanged, in the downloadable reproducibility bundle as a documented superseded reference.

**11**. Priority miss penalty and study size boost corrected (v3.0). The 0.85× priority-miss penalty (applied when a platform scores 1/5 on a dimension the user has prioritised) had been gated on the context-boosted effective weight rather than the user's own slider value, so an automatic goal- or matrix-based adjustment could trigger a penalty on a dimension the user never touched; it now gates on the raw slider. Separately, the large/extra-large study-size boost to cost and throughput (previously +1/+2) had applied the same input at full strength to both dimensions simultaneously, dimensions that happen to be Nomic nELISA's strongest, tied-max scores; the boost is now halved (+0.5/+1).

**12.** Sample-flexibility weight and priority-miss extension corrected (v3.0). The tissue sample-flexibility weight (8) was disproportionate against the same weight for cell culture and multiple sample types (5 each), with no documented reason tissue should carry more: at 8, a platform with the maximum flexibility score earned more from matrix compatibility alone (40 points) than the maximum possible contribution from a fully-maxed priority slider (25 points), letting sample type override any explicit user priority. Tissue now matches the other two atypical matrices at 5; the underlying platform-specific overrides, reflecting genuine validation differences, are unchanged. This alone left a related gap open: a platform with no published validation for the selected sample type could still win outright on strength in unrelated dimensions. Before the correction, Nomic nELISA, which has no published solid-tissue validation, won 3.28% of tissue-matrix scenarios outright on cost and throughput strength alone; after it, its tissue outright rate is 0%. The priority-miss penalty (0.85×, described in step 11) now also fires when a platform's sample-flexibility score for the selected matrix is 1/5 (no published validation at all), closing this gap. None of the corrections in steps 10 through 12 altered any platform score; all are corrections to how user inputs translate into effective weights or how the simulation tallies its results.

**13.** Biognosys cost score corrected and tissue override removed (v3.5). An evidence review of commercial pricing placed all four broad discovery platforms (Olink Explore HT, SomaSeq, Seer Proteograph XT, and Biognosys TrueDiscovery) in the same price band at volume, so Biognosys TrueDiscovery's cost-per-sample score rose from 2 to 3; the previous score rested on older service quotes that no longer reflect current pricing. The step 7 tissue override on Olink Explore HT and SomaSeq was removed: homogenate and lysate input is the standard way affinity assays are run on tissue, both platform lineages have published tissue applications, and penalising them against a native solid-tissue MS criterion conflated technology class with validation. Both platforms revert to their standing multi-matrix score of 3 in tissue; platforms with no published solid-tissue validation retain 1. Net effect on the baseline: Biognosys TrueDiscovery's co-recommendation roughly doubles (8.1% to 16.5%) while its outright rate falls (1.4% to 1.0%), and tissue scenarios shift from a single dominant platform to a contested three-way space (Biognosys 17.8%, SomaSeq 13.0%, Olink 3.2% outright). All values in this manuscript reflect this configuration (Supplementary Table S10).

The fairness simulation incorporates the CNS-focus and longitudinal/repeat-sampling toggles as a 2 × 2 factorial but excludes the absolute quantification and PTM detection toggles; under v4.0 an absolute-quantification requirement is instead resolved by the capability filter described in the main methods (only 7 of the 65 recorded responses set it). All four axes, goal, sample type, study size, and toggle state, are now reweighted by observed real-world prevalence rather than treated as equally likely (CNS-focus 7.7%; longitudinal toggle imputed at approximately 17.5%; Supplementary Table S10), replacing an earlier approach that reweighted only the two toggles and left the other axes uniform, itself retained as a superseded reference (Supplementary Table S11); the equal-sample-type version this superseded had slightly overrepresented tissue and cell culture relative to real-world usage, where plasma and serum dominate, a distortion the current prevalence weighting corrects directly rather than merely disclosing. Platform scores reflected published literature as of the manuscript submission date and will require updating as new validation data emerge.

### Supplementary Note S2. Technical implementation

We developed APT using React 19 with Vite as the build system and Tailwind CSS v4 for styling. We compiled the application to a self-contained single HTML file using vite-plugin-singlefile, eliminating external runtime dependencies and enabling deployment to static hosting without a server-side component. We maintained platform metadata and scoring data as JSON configuration files under version control, allowing independent score updates without changes to application logic. We deployed the application via Vercel at https://aptatlas.org/. We make the source code available at https://github.com/christopherdwhelan/aptatlas/.

The tool included the following interactive features: a weighted recommendation tool with real-time scoring updates; a platform comparison view with radar charts and tabular scoring breakdowns; a curated evidence database of published head-to-head studies; a protein target lookup enabling platform selection based on coverage of user-specified gene lists; and this methods documentation.

### Supplementary figure legends

**Supplementary Figure S1.** Per-tool fairness breakdown by the development version that originally served each cohort. (A) Profiles served by earlier tool versions (v1.0 to v1.2.2; 47 served, 45 scored): outright top-pick and top-tier co-recommendation shares per platform. (B) Profiles served by later versions (v2.0 to v2.2; 18 served and scored): the same two measures. Two panel-A profiles requiring systematic PTM detection are excluded by the capability filter. Cohorts were assigned by joining response timestamps to the public change log. Tie rates were 64% (median first-to-second gap 2 points) for the earlier-versions cohort and 67% (median gap 2 points) for the later-version cohort. Both panels re-score the recorded inputs through the final production engine to permit a like-for-like comparison and are not the recommendations users observed at the time.

**Supplementary Figure S2.** Ordinal platform scores grounded in documented values. For four dimensions, the documented evidence for each platform is shown beside its assigned score (1 to 5). (A) Proteome coverage: distinct proteins or protein groups; hatched extensions mark documented larger values from enhanced workflows or newer product versions (Seer Proteograph ONE; Biognosys P2-enriched; NULISA panel configurations). (B) Measurement precision: median coefficient of variation, one point per published study; Nomic Omni is shown as its published bound (inter-assay CVs predominantly below 15%, Dagher et al., 2025); the NULISA value is the median intraplate CV from a 92-target head-to-head against Olink Explore (Feng et al., 2023). (C) Sensitivity: the pg/mL band of each assigned score under the scoring rubric (Supplementary Table S2); the arrow denotes the open-ended highest band (<0.01 pg/mL). (D) Cost: estimated service price per sample. These are informal estimates assembled from public price lists and quotes received by the authors, not published values, and boundaries are indicative; solid bars for Olink Explore HT and SomaSeq reflect published price-list ranges, truncated at a common $800 ceiling; leftward arrows denote estimated large-volume discounting below the shown floor; Nomic Omni's $50 is a vendor-published list price.

**Supplementary Figure S3.** Worked examples of how study context maps to a recommendation. Six representative scenarios and the platform Help Me Choose recommends for each, with the percentage of matched scenarios the platform leads: (A) population pQTL mapping (Seer

Proteograph XT, outright in 100% of matched scenarios, the most decisive single-dimension result in the simulation), (B) CNS biomarker discovery in CSF (Alamar NULISA, outright in 34.5%; Olink Explore HT leads co-recommendation at 50.8%), (C) large plasma biomarker discovery (Seer Proteograph XT, co-recommended at 45.8%, with Olink Explore HT co-recommended at 38.9%), (D) a study requiring absolute concentrations (Nomic Omni, the sole platform passing the capability filter, recommended in 100% of matched scenarios), (E) tissue proteomics (Biognosys TrueDiscovery, co-recommended in 63.4% and outright in 17.8%; SomaSeq and Olink Explore HT win 13.0% and 3.2% outright), and (F) a population-scale screen (Nomic Omni, co-recommended in 46.5%, with Seer Proteograph XT at 31.5%). Each scenario conditions on its named study context and sweeps all remaining inputs uniformly; recommendations are production version outputs (Figure 4A). Driver dimensions follow the goal weight map in Supplementary Table S11.

**Supplementary Figure S4.** Coverage of druggable target classes across six plasma proteomics platforms. Horizontal bars show the percentage of each protein class's druggable targets detectable by the union of all six platforms, ordered from best to least covered and coloured by family skew (affinity-skewed, mass-spectrometry-skewed, or balanced, where skew denotes an at least 15 percentage-point gap between any-affinity and any-mass-spectrometry coverage). Overlaid markers show the fraction reachable by any affinity platform (circle) and any mass spectrometry platform (square). Ion channels (30.8% of 156 targets) and G-protein-coupled receptors (47.4% of 154) are the least-covered classes, while transporters are the single class where mass spectrometry leads affinity panels. Underlying values are in Data S2 (Supplementary Table S14).

**Supplementary Figure S5.** Simulated against real-world outright-win rate per platform. Outright wins are cases where a single platform is recommended with no co-recommended tie, the counterpart to the co-recommended comparison in Figure 4C. Blue bars give the prevalence-weighted simulated baseline across 1,259,712 enumerated scenarios; orange bars give the observed rate across the 63 recorded researcher profiles, with counts shown because one profile is 1.6% of that cohort. Four of the six platforms take no outright win in the observed cohort. This is a property of the tie logic rather than of those platforms: most real profiles resolve to co-recommended ties, where two or more platforms fall within the tool's 3-point tie band and none is reported as a sole winner, so the outright statistic excludes the majority of real recommendations by construction. Figure 4C reports the co-recommended rate for the same profiles and is the statistic that reflects what most users see.

**Supplementary Figure S6.** Fairness of the Help Me Combine recommendation across the space of possible user priorities. (A) Win-share by pair for all 15 pairs, from the deterministic priority sweep of 231 combinations of one to three of the eleven priorities evaluated against four coverage denominators, 924 scenarios in total. The dashed line marks the share expected under uniform pair selection (100% / 15 = 6.7%). No pair exceeds 18.7% of scenarios and 14 of 15 win at least one. (B) Platform participation: the percentage of scenarios in which each platform appears in the winning pair. Because every scenario returns one pair of two platforms, the six shares sum to 200%, and the dashed line marks the 33.3% each platform would take under even participation. Nomic Omni and Seer Proteograph XT participate most, each in roughly half of winning pairs, and Alamar NULISA's lower participation reflects its narrower panel of approximately 385 assays, which wins where its strengths in precision and sensitivity are prioritised. Underlying values are in Supplementary Table S12.
