## Supplementary figures and images for "Atlas of Proteomic Technologies: an evidence-based framework for selecting and combining commercial proteomics platforms"

### Figure S2

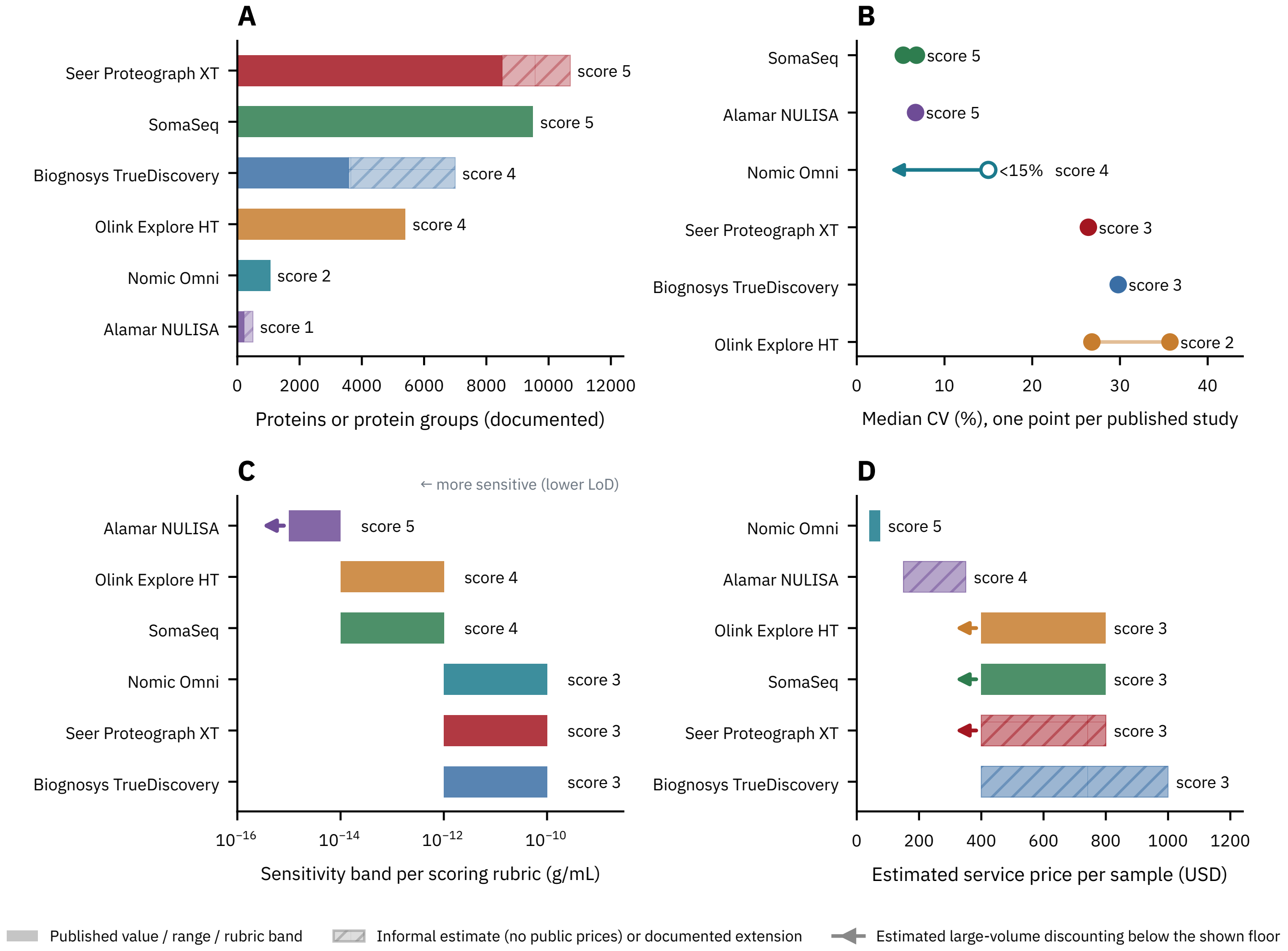

### Figure S6

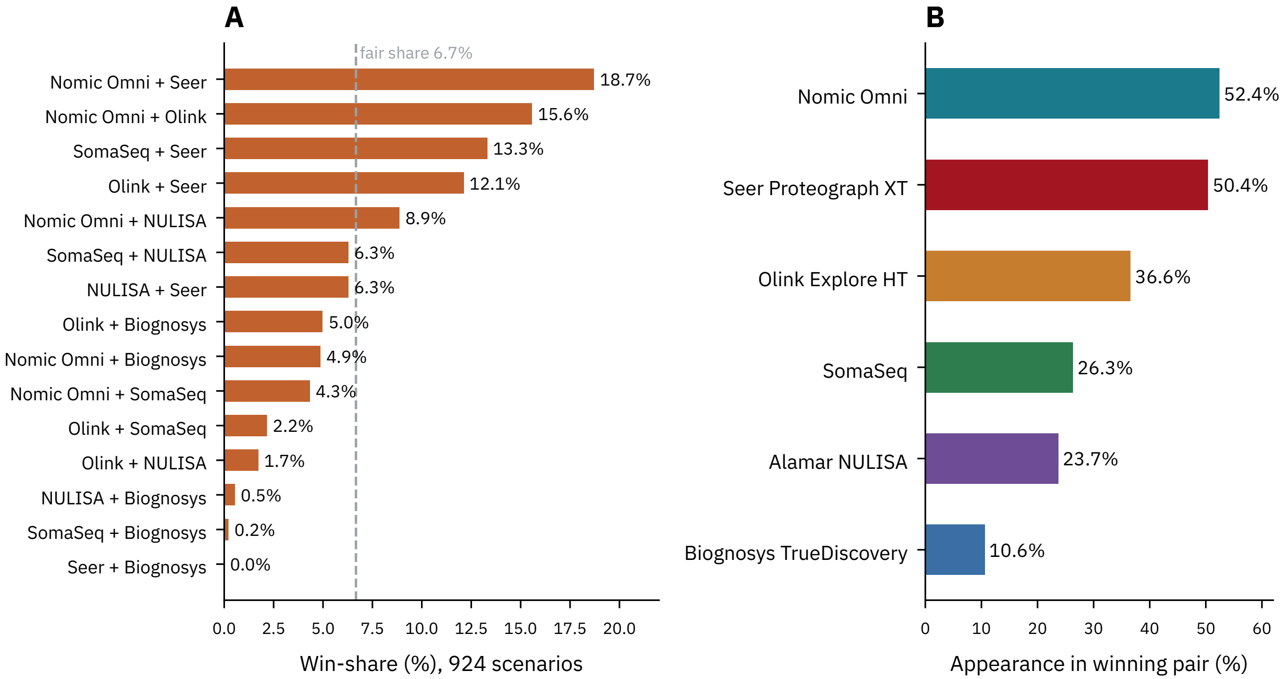
