## Supplementary material for "Atlas of Proteomic Technologies: an evidence-based framework for selecting and combining commercial proteomics platforms": Figure S5

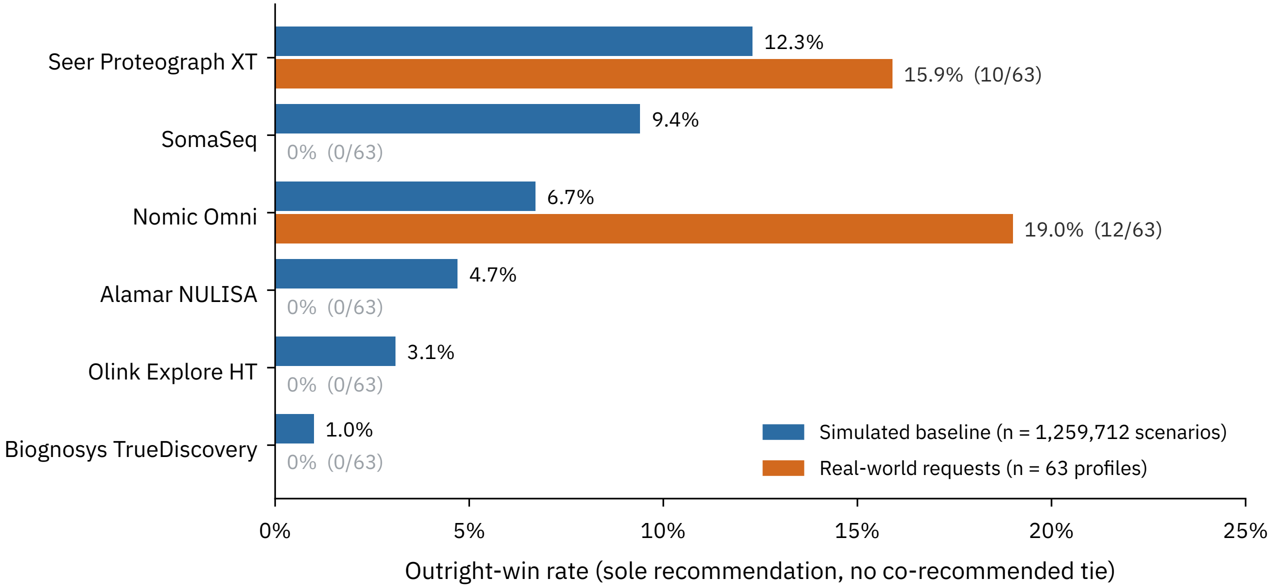


One real-world profile = 1.6%. Four platforms take no outright win in the observed cohort:
most real profiles resolve to co-recommended ties (Figure 4C), which the outright statistic excludes.
