## Supplementary material for "Atlas of Proteomic Technologies: an evidence-based framework for selecting and combining commercial proteomics platforms": Figure S4

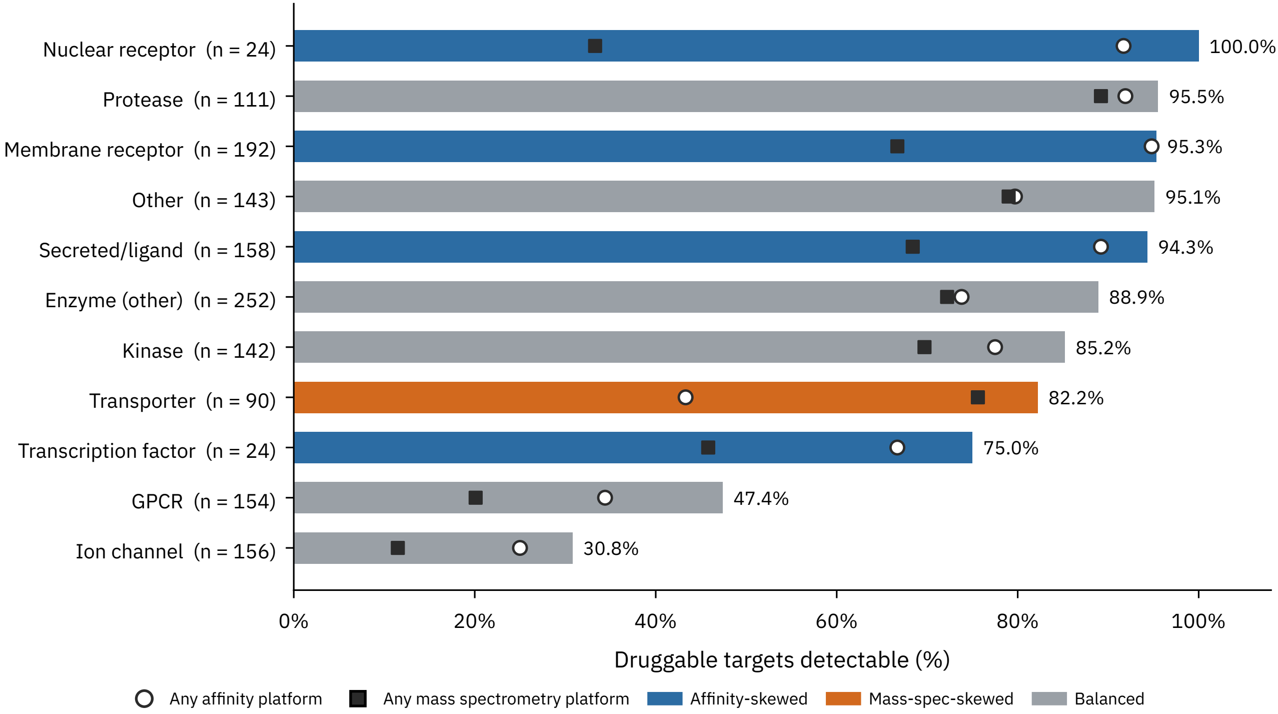


Bars give coverage by the union of all six platforms. Skew denotes a gap of at least 15 percentage points between any-affinity and any-mass-spectrometry coverage.
