## Supplementary material for "Atlas of Proteomic Technologies: an evidence-based framework for selecting and combining commercial proteomics platforms": Figure S3

### A Population pQTL mapping

pQTL slider maxed - plasma, all goals and sizes

**Seer Proteograph XT**

100.0% of matched scenarios, outright

pQTL accuracy 5/5; the most decisive single-dimension result

### B CNS biomarker discovery

discovery goal - CSF - CNS focus on

**Alamar NULISA**

34.5% of matched scenarios, outright

CNS sensitivity boost; Olink leads co-recommendation (50.8%)

### C Large biomarker discovery

discovery goal - plasma - 1,000-10,000 samples

**Seer Proteograph XT**

45.8% of matched scenarios, co-recommended

Coverage-led; Olink Explore HT co-recommended (38.9%)

### D Absolute concentrations required

discovery - plasma - absolute-quantification filter on

**Nomic Omni**

100.0% of matched scenarios, outright

Sole platform passing the capability filter (pg/mL output)

### E Tissue proteomics

tissue matrix - all goals and sizes

**Biognosys TrueDiscovery**

63.4% of matched scenarios, co-recommended

Multi-matrix 5/5; SomaSeq (13.0%) and Olink (3.2%) win some scenarios outright

### F Population-scale screen

population goal - plasma - >10,000 samples

**Nomic Omni**

46.5% of matched scenarios, co-recommended

Cost and throughput boosts, both Nomic 5/5; Seer co-recommended (31.5%)
