## Supplementary material for "Atlas of Proteomic Technologies: an evidence-based framework for selecting and combining commercial proteomics platforms": Figure S1

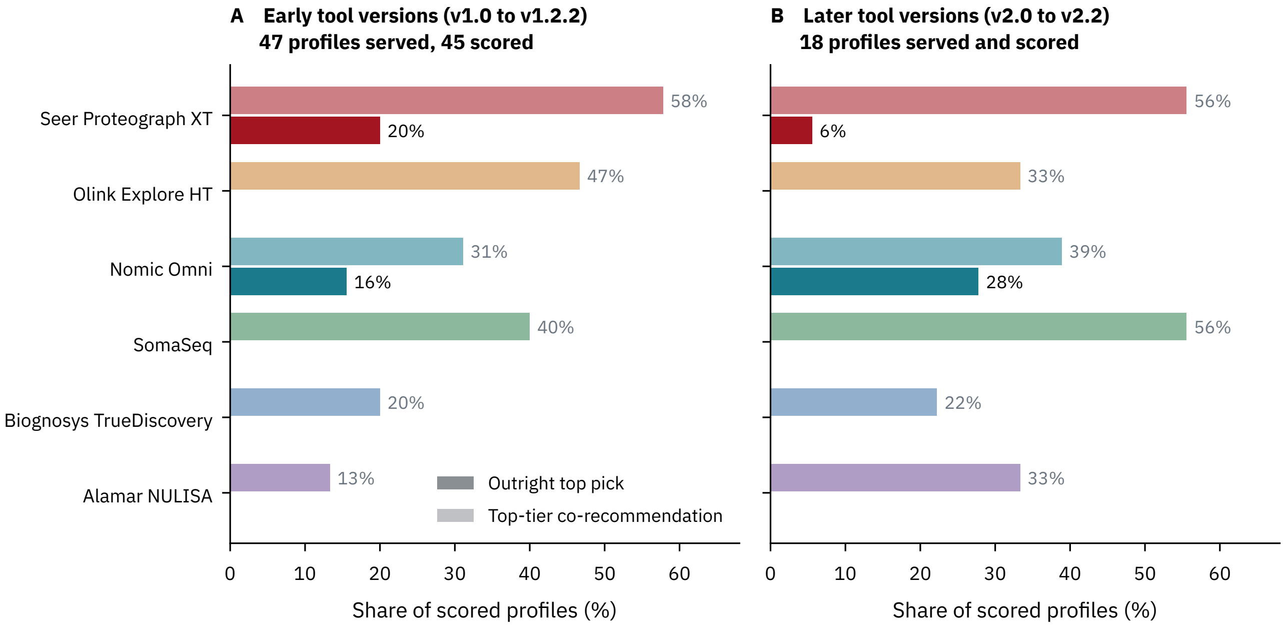


All profiles re-scored through the final production engine (like-for-like); not the recommendations served at the time.
Version assignment from response timestamps joined to the public change log. Two early-cohort profiles requiring systematic PTM detection are excluded by the capability filter. Tie rates: 64% (early) and 67% (later); median first-to-second gap 2 points in both.
